# Insulin determines the effects of TGF-β on HNF4α transcription and epithelial-to-mesenchymal transition in hepatocytes

**DOI:** 10.1101/2023.01.12.523351

**Authors:** Rilu Feng, Chenhao Tong, Tao Lin, Hui Liu, Chen Shao, Yujia Li, Carsten Sticht, Kejia Kan, Xiaofeng Li, Rui Liu, Sai Wang, Shanshan Wang, Stefan Munker, Hanno Niess, Christoph Meyer, Roman Liebe, Matthias P Ebert, Steven Dooley, Hua Wang, Huiguo Ding, Hong-Lei Weng

**Author notes:** Corresponding author: Hong-Lei Weng, Department of Medicine II, Section Molecular Hepatology, University Medical Center Mannheim, Medical Faculty Mannheim, Heidelberg University Theodor-Kutzer Ufer 1-3, 68167 Mannheim, Germany. The authors contributed equally and share first authorship (Rilu Feng, Chenhao Tong).

## Abstract

To date, epithelial-to-mesenchymal transition (EMT) has been observed in cultured hepatocytes, but not *in vivo*. TGF-β is supposed to initiate EMT in hepatocytes by inhibiting HNF4*α* through the SMAD2/3 complex. We report that TGF-*β* does not directly inhibit HNF4*α*, but contributes to its transcriptional regulation by SMAD2/3 recruiting acetyltransferase CBP/p300 to the HNF4*α* promoter. The recruitment of CBP/p300 is indispensable for C/EBP*a* binding, another essential requirement for constitutive HNF4*α* expression in hepatocytes. In contrast to the observed induction of HNF4*α*, SMAD2/3 inhibits C/EBP*α* transcription. Therefore, long-term TGF-*β* incubation results in C/EBP*α* depletion, which abrogates HNF4*α* expression. Intriguingly, SMAD2/3 inhibitory binding to the C/EBP*α* promoter is abolished by insulin. Thus, maintaining a high insulin concentration in culture medium ensures constitutive HNF4*α* and thereby prevents TGF-β-induced hepatocyte EMT. Furthermore, insulin inhibits TGF-*β*-induced SMAD2/3 binding to the promoters of core EMT transcription factors e.g., SNAI1. SNAI1 transcription requires both SMAD2/3 and FOXO1 in nuclei. Insulin inhibits SNAI1 transcription through impeding SMAD2/3 binding to its promoter and inducing FOXO1 phosphorylation. Hence, insulin is the key factor that prevents TGF-*β*-induced EMT in hepatocytes.

## Introduction

The role of epithelial-to-mesenchymal transition (EMT) in organ fibrosis (including in the liver, lung, and kidney) is a long-term subject of active debate^1^. It has been shown that cultured hepatocytes, alveolar or renal epithelial cells differentiate into a myofibroblast-like phenotype under stimulation with transforming growth factor (TGF)-*β*^1^. However, *in vivo* lineage-tracing analyses have not provided compelling evidence to date that fibrosis-associated myofibroblasts are derived from epithelial cells^1^. In the liver, hepatocyte nuclear factor 4*α* (HNF4*α*) is a master hepatic transcription factor to regulate key hepatocyte functional genes and maintain epithelial identity^2–5^. In adult hepatocytes, HNF4*α* regulates more than 40% of all functional genes through binding to their promoters^6^. HNF4*α*-regulated genes are crucial in numerous physiological functions, including synthesis of coagulation factors, lipid transport, fatty acid oxidation, bile acid synthesis and transport, and metabolism of glucose, lipoproteins, steroids and amino acids^7^. Low expression of HNF4*α* is implicated in a series of liver diseases such as metabolic syndrome, diabetes, cholestasis, cirrhosis and hepatocellular carcinoma (HCC)^7, 8^. In patients with acute liver failure (ALF), loss of hepatic HNF4*α* expression is closely associated with poor prognosis^9^. These observations highlight a key role of HNF4*α* in controlling liver function and maintaining epithelial identity^2–5, 10^.

Santangelo and colleagues showed that in cooperation with HNF1*α*, HNF4*α* inhibited key EMT master regulator genes in both cultured hepatocytes and animals, and a loss of HNF4*α* leads to EMT-like alterations in hepatocytes^11^. In diseased livers, inflammation is the main factor inhibiting hepatic HNF4*α* expression^12^. Among a plethora of inflammatory factors, TGF-*β* is the best known HNF4*α* inhibitory cytokine^13^. In cultured hepatocytes, TGF-*β* inhibits HNF4*α* expression by upregulating snail family transcriptional repressor (SNAI), a transcription repressor that binds to the *Hnf4a* promoter, leading to EMT of hepatocytes^14^. However, high levels of TGF-*β* do not always lead to loss of HNF4*α* in hepatocytes. As in other organs, lineage-tracing analyses deny the occurrence of hepatocyte EMT in animal models^15^. A previous study had examined serum TGF-*β* concentrations and liver p-SMAD2 expression in chronic liver disease patients with different disease severity^16^. TGF-*β* and p-SMAD2 levels positively correlated with inflammatory grades and fibrotic stages^16^. However, HNF4*α* loss only occurred in a portion of cirrhotic patients^9^. These observations raise two interesting questions: What is the underlying reason that hepatocytes in patients with chronic liver disease are capable of maintaining HNF4*α* expression in spite of high levels of TGF-*β*, while hepatocytes in other patients lose HNF4*α* expression when their disease progresses into liver cirrhosis or terminal liver failure? In addition, why do hepatocytes undergo EMT only in tissue culture, but not *in vivo*?

In this study, we find that HNF4*a* transcription in hepatocytes requires binding of both SMAD2/3 and C/EBP*a* to its promoter. Although SMAD2/3 binding does not directly influence *HNF4A* transcriptional activity, SMAD2/3-recruited acetyltransferase CBP/p300 is essential for C/EBP*a* binding. Interestingly, SMAD2/3 simultaneously represses C/EBP*a* transcription. Physiologically, insulin maintains C/EBP*a* expression by inhibiting SMAD2/3 binding to its *CEBPA* promoter. In the condition of severe inflammation, tumor necrosis factor alpha (TNF-*a*)-induced insulin resistance in hepatocytes results in TGF-*b*-induced SMAD2/3 inhibition of C/EBP*a* expression. The ensuing depletion of C/EBP*a* abrogates HNF4*a* expression in hepatocytes. Given that the half-life of insulin is approximately 1 hour, depletion of insulin over time results in hepatocytes undergoing EMT *in vitro*. Continuously replenishing insulin in culture medium keeps SMAD2/3 away from to the *CEBPA* promotor, and thus prevents hepatocytes from TGF-*b*-induced EMT.

## Results

### In normal hepatocytes, constitutive HNF4*α* expression requires TAF6/9 and H3K4me3

To clarify how hepatocytes constitutively express HNF4*α* (**Figure 1A**), we first scrutinized the binding of RNA polymerase II (Pol II) to the core promoter (defined as ±50bp around the transcription start site, TSS) of the *HNF4A* gene^17^. In human primary hepatocytes (HPHs) and mouse hepatocyte line AML12 cells, chromatin immunoprecipitation (ChIP) assays showed that the *HNF4A* core promoter was bound by Pol II with phosphorylation on serine 5 (S5) and serine 2 (S2) of the heptapeptide repeats in the C-terminal domain of the Rbp1 subunit (**Figure 1B**), indicating transcription initiation and elongation of this gene. DNA sequence analysis revealed that the *HNF4A* core promoter does not possess a classical TATA box, but has a downstream promoter element (DPE) (**Figure 1C**). ChIP assay shows that the *HNF4A* core promoter is bound by histone 3 (**Figure 1B**). Given that transcription factor II D binding to the core promoter of genes without TATA box usually requires H3K4me3- marked nucleosomes^18^, we examined H3K4me3 binding in the *HNF4A* core promoter. ChIP assay shows that the *HNF4A* core promoter did indeed bind H3K4me3-marked nucleosomes in HPHs and AML12 cells (**Figure 1B**). Inhibition of H3K4me3 with lysine-specific demethylase 5C significantly reduced mRNA and protein expression of HNF4*α* in AML12 cells and mouse primary hepatocytes (MPHs, **Figure 1D-E**).

**Figure 1.**
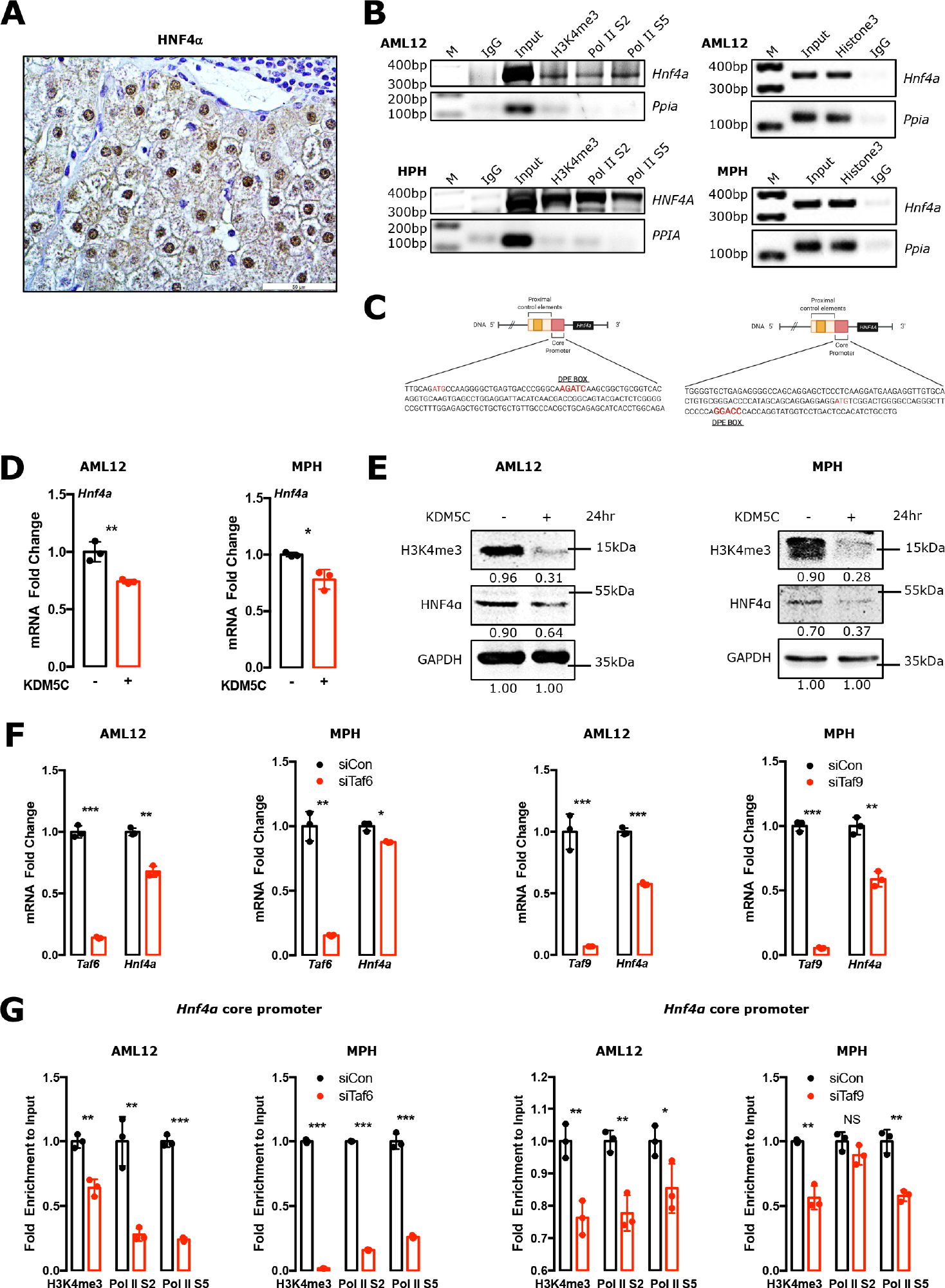
Constitutive HNF4*α* expression in hepatocytes requires TAF6, TAF9 and H3K4me3. **(A)** Immunohistochemical staining shows HNF4*α* expression in normal liver tissue. Original magnification: x40. **(B)** Chromatin immunoprecipitation (ChIP) assays examined H3K4me3, RNA polymerase II phosphorylation of Ser2 (Pol II S2) and Pol II S5 binding to the *HNF4A* core promoter (AML12: -172∼+160base pair (bp), human primary hepatocytes (HPHs): -250∼+67bp), and Histone 3 binding to the *Hnf4a* promoter (-172∼+160bp) in AML12 cells and mouse primary hepatocytes (MPHs). **(C)** A scheme depicts the *HNF4A* core promoter sequence and downstream promoter element (DPE) box in human and mouse species. **(D-E)** qPCR **(D)** and Western blotting **(E)** analyzed HNF4*α* expression in AML12 and MPHs treated with 3nM Lysine- specific demethylase 5C (KDM5C) for 24 hours. **(F)** qPCR analyzed HNF4*α* expression in AML12 cells and MPHs transfected with TBP-associated factors 6 or 9 (Taf6 or Taf9) small interfering RNA (siRNA) for 24 hours. **(G)** ChIP-qPCR examined the binding activities of H3K4me3, Pol II S2 and Pol II S5 on the *Hnf4a* promoter (- 172∼+160bp) in AML12 cells and MPHs treated with *Taf6* or *Taf9* siRNA for 24 hours. Bars represent the mean ± SD, *P*-values were calculated by unpaired Student’s t test, *, *P*<0.05; **, *P*<0.01; ***, *P*<0.001; and NS, No significance. Triple experiments were performed, and one representative result is shown.

Given that the DPE motif is bound by TBP-associated factor 6 (TAF6) or TAF9 subunit of transcription factor II D^19^, we knocked down *Taf6* and *Taf9* by RNA interference (RNAi) in AML12 cells and MPHs (**Figure 1F**). Quantitative polymerase chain reaction (qPCR) assay showed that mRNA expression of *Hnf4a* was significantly reduced when *Taf6* or *Taf9* was knocked down (**Figure 1F**). ChIP assays further revealed that knockdown of *Taf6* or *Taf9* decreased both S5-P and S2-P Pol II binding to the *Hnf4a* core promoter (**Figure 1G**), indicating the effect of TAF9 and TAF6 on *Hnf4a* transcription.

### Constitutive *HNF4A* transcription requires transcription regulators SMADs and C/EBP*α* binding to the promoter

Sufficient transcription requires transcription activators that bind to the respective gene promoter. To clarify the transcriptional activators that bind to the *Hnf4a* promoter, we analyzed the *Hnf4a* gene regulatory sequences. Transcription regulator C/EBP*α* possesses binding sites on the *Hnf4a* promoter (**Figure 2A**). In HepG2 cells, ChIP sequencing analyses showed SMAD3 binding to the *HNF4A* promoter (**Figure 2B**). Subsequently, ChIP assays confirmed SMAD2, SMAD3 and C/EBP*α* binding to the *Hnf4a* promoter in AML12 cells (**Figure 2C**). Knockdown of *Smad2*, *Smad3* and *Cebpa* by RNAi inhibited Pol II binding to the *Hnf4a* core promoter (**Figure 2D**) and mRNA and protein expression of HNF4*α* in hepatocytes (**Figure 2E-F**). Ectopic expression of *Cebpa* robustly increased mRNA and protein expression of HNF4*α* in hepatocytes (**Figure 2E-F**).

**Figure 2.**
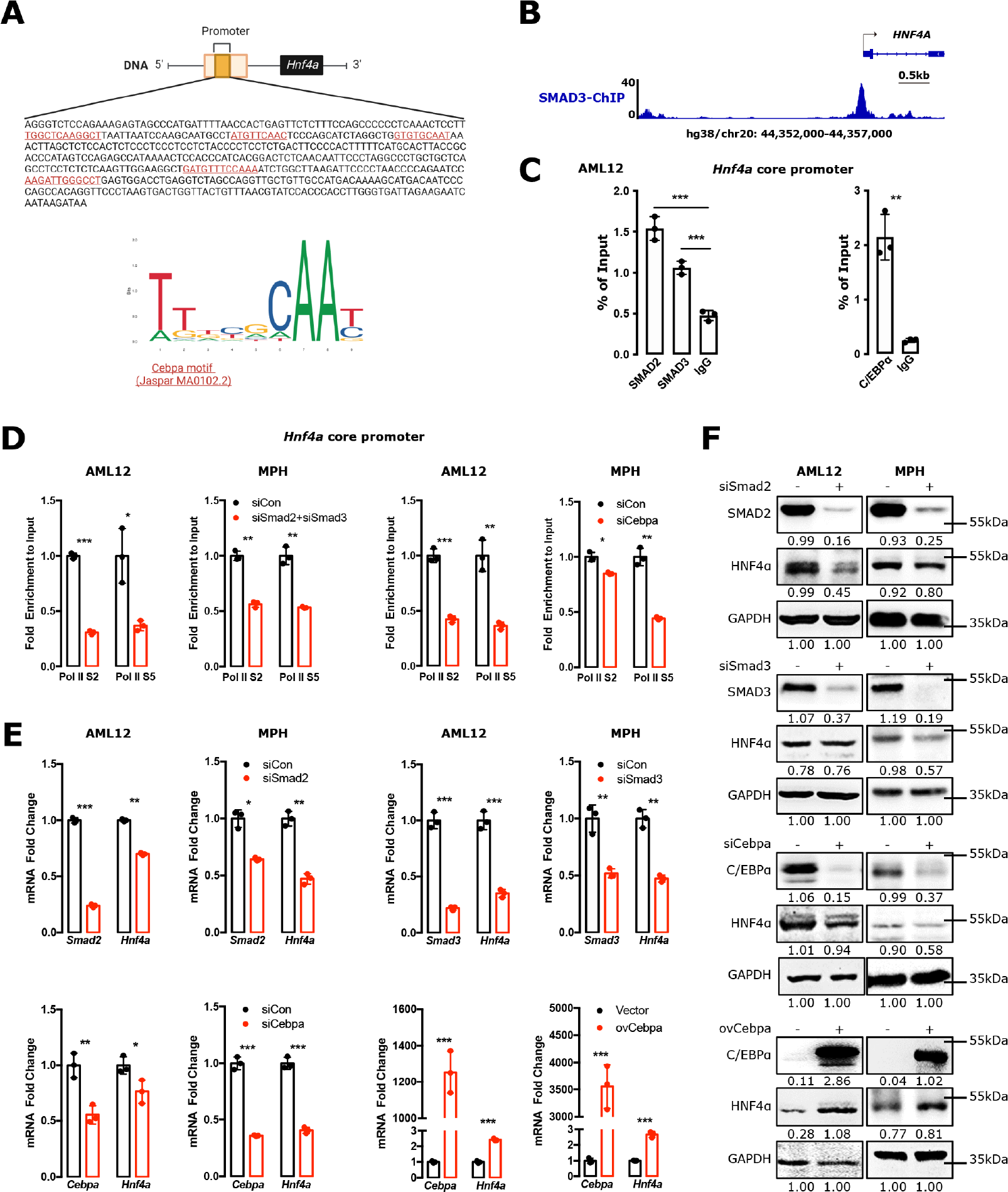
Constitutive *HNF4A* transcription requires transcription regulators SMADs and C/EBP*α* binding to the promoter. **(A)** The binding sites of C/EBP*α* on the *Hnf4a* promoter (-500∼0bp) was predicted by Jaspar dataset^46^. **(B)** ChIP sequencing showed enrichment profiles of SMAD3 on *HNF4A* promoter in HepG2 cells. **(C)** ChIP-qPCR examined SMAD2, SMAD3 and C/EBP*α* binding to the *Hnf4a* promoter (-350∼0bp) in AML12 cells. **(D)** ChIP-qPCR measured the binding activities of Pol II S2 and Pol II S5 on the *Hnf4a* core promoter (-172∼+160bp) in AML12 cells and MPHs transfected with *Smad2*, *Smad3* or *Cebpa* siRNA for 24 hours. **(E-F)** qPCR **(E)** and Western blotting **(F)** analyzed the effects of SMAD2, SMAD3 and C/EBP*α* on HNF4*α* expression in AML12 cells and MPHs transfected with *Smad2*, *Smad3* or *Cebpa* siRNA for 24 hours, or *Cebpa* construct for 48 hours. Bars represent the mean ± SD, *P* values were calculated by unpaired Student’s t test, *, *P*<0.05; **, *P*<0.01, and ***, *P*<0.001. Triple experiments were performed, and one representative result is shown.

### C/EBP*α* binding to the *HNF4A* promoter requires SMAD2/3-recruited CBP/p300

Impressively, co-immunoprecipitation analysis showed that SMAD2 and SMAD3 did not interact with C/EBP*α* in hepatocytes (**Figure 3A**). qPCR and Western blotting analyses showed that forced overexpression of *CEBPA* was not capable of restoring mRNA and protein expression of HNF4*α* when SMAD2/3 were reduced (**Figure 3B- C**). To confirm the role of SMAD and C/EBP*α* in *Hnf4a* transcription, we established a *Hnf4a* luciferase construct and transfected them into HEK293 cells. Overexpressing *Cebpa*, but not *Smad2* or *Smad3*, remarkably increased *Hnf4a* luciferase activity (**Figure 3D**). These results suggest that C/EBP*α* binding to the *HNF4A* promoter directly regulates gene transcription. Although SMAD2/3 binding to the *HNF4A* promoter does not increase gene transcription, it is required for C/EBP*α* binding.

**Figure 3.**
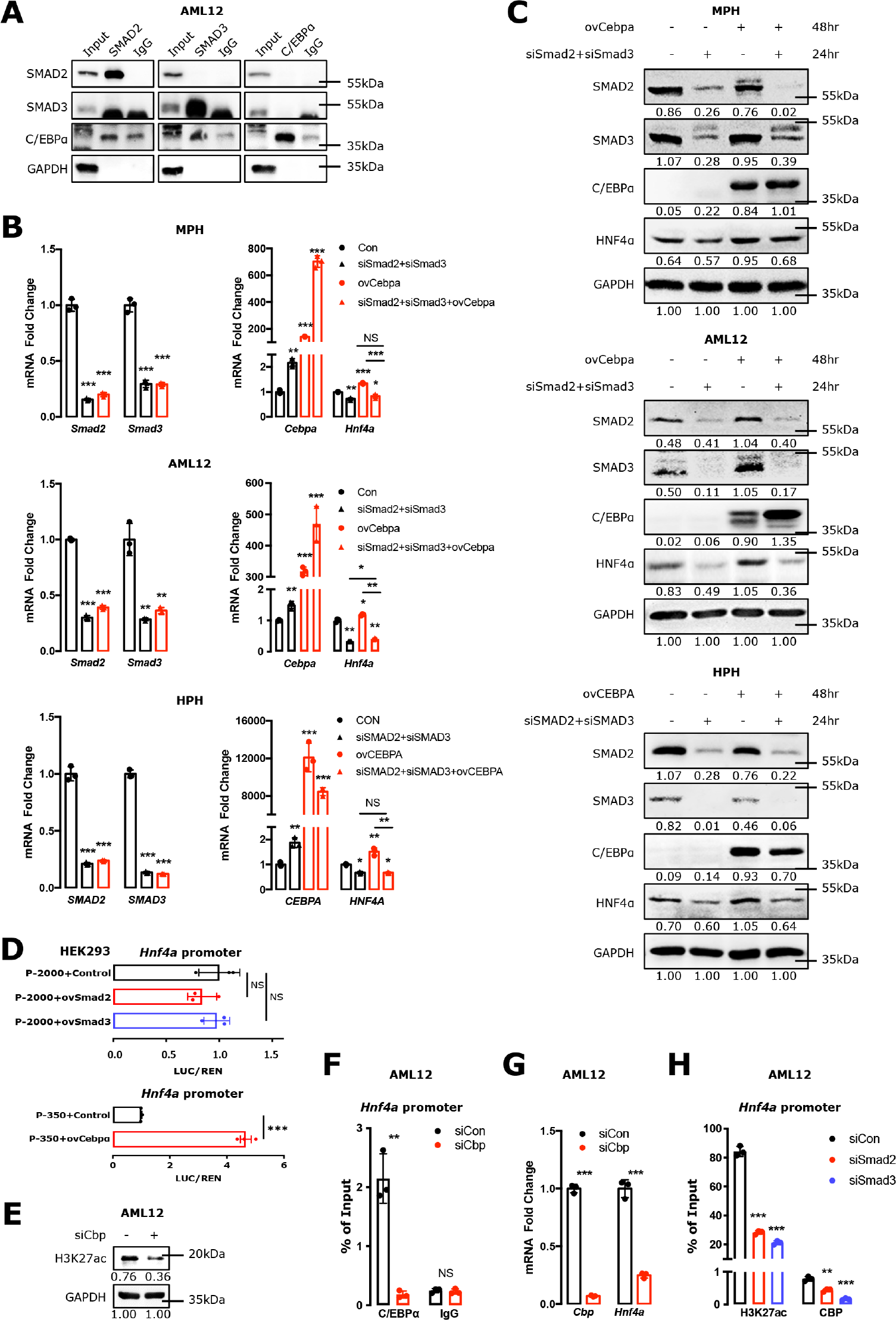
C/EBP*α* binding to the *HNF4A* promoter requires SMAD2/3-recruited CBP/p300 **(A)** Co-immunoprecipitation examined the protein interactions between SMADs and C/EBP*α* in AML12 cells. **(B-C)** qPCR **(B)** and Western blotting **(C)** analyzed HNF4*α* expression in AML12 cells, MPHs and HPHs transfected with *SMAD2*, *SMAD3* siRNA for 24 hours, followed by *CEBPA* construct transfection for 48 hours. **(D)** Luciferase reporter assay analyzed the activity of *Hnf4a* promoter (-350∼0bp, -2000∼0bp) in HEK293 cell transfected with or without *Cebpa*, *Smad2*, *Smad3* plasmid, respectively. **(E)** Western blotting examined H3K27ac expression; **(F)** ChIP-qPCR analyzed C/EBP*α* binding activity on *Hnf4a* promoter (-350∼0bp); **(G)** qPCR showed HNF4*α* expression. The cells were treated with *Cbp* siRNA for 24 hours. **(H)** ChIP-qPCR examined H3K27ac and CBP binding activities on *Hnf4a* promoter (-214∼-58bp) in AML12 cells transfected with *siSmad2* or *siSmad3* for 48 hours. Bars represent the mean ± SD, *P*-values were calculated by unpaired Student’s t test and One-way ANOVA, *: *P*<0.05; **: *P*<0.01; ***: *P*<0.001; and NS, no significance. Triple experiments were performed, and one representative result is shown.

How does SMAD2/3 recruitment to the *HNF4A* promoter influence C/EBP*α*- dependent *HNF4A* transcription? It is commonly known that the presence of the histone acetyltransferase CREB-binding protein (CBP)/p300 is required for gene transcription^20^. Binding of CBP/p300 to SMAD complex has been shown in different cells^21^. Therefore, we speculated that SMADs might be essential for CBP/p300 recruitment to the *Hnf4a* promoter. As expected, knockdown of *Cbp/p300* by RNAi remarkably reduced H3K27ac expression, C/EBP*α* binding to the *Hnf4a* promoter, and thus inhibited mRNA expression of *Hnf4a* (**Figure 3E-G**). Subsequently, we knocked down SMAD2 and SMAD3 by RNAi (**Figure 3H**). In the absence of SMAD2/3, both CBP/p300 and H3K27ac binding to the *Hnf4a* promoter was prevented (**Figure 3H**).

These results clarify the distinct effects of SMAD proteins and C/EBP*α* in the regulation of *Hnf4a* transcription.

Given that SMAD proteins are localised in the cytoplasm of cells, and that only activated SMADs efficiently enter the nucleus of cells to act as transcription factors^22, 23^, the above results raised a question: whether non-activated SMAD proteins were able to enter nuclei and initiate gene transcription in normal liver or cultured hepatocytes. We performed IHC for p-SMAD2 in 20 normal mouse liver tissues and observed that hepatocytes in normal livers expressed p-SMAD2 (**Figure S1A**). In cultured AML12 cells, SB431542, a TGF-*β* receptor I inhibitor, significantly inhibited p-SMAD2 and p-SMAD3 expression (**Figure S1B**). In these hepatocytes without external TGF-*β* stimulation, SB431542 remarkably reduced SMAD2/3 binding to the promoters of the *Hnf4a* and *Cebpa* gene (**Figure S1C**). These results reveal the existence of basic TGF-*β* levels *in vivo* and *in vitro*.

### Mediator complex is required for sufficient *HNF4A* transcription

To sufficiently initiate gene transcription, a substantial portion of genes require the Mediator protein complex, which links transcriptional activator bound promoter and enhancer to general transcription factors^24^. Therefore, we examined the effects of Mediator14 (MED14) and Cyclin dependent kinase 8 (CDK8), two components of the Mediator complex, on basal *HNF4A* transcription. In AML12 cells and MPHs, knockdown of *Med14* reduced mRNA and protein expression of HNF4*α* (**Figure 4A, B**), but did not alter C/EBP*α* expression (**Figure 4C**). Furthermore, knocking down *Med14* reduced both S5-P Pol II and S2-P Pol II binding to the *Hnf4a* core promoter, as well as SMAD2/3 and C/EBP*α* binding to the *Hnf4a* promoter (**Figure 4D**). Like *Med14*, knocking down *Cdk8* by RNAi significantly reduced mRNA expression of *Hnf4a* in AML12 cells and MPHs (**Figure 4E**). CDK8 reduction did not alter *Cebpa* mRNA expression in MPHs, but decreased *Cebpa* mRNA expression in AML12 cells (**Figure 4F**). These results suggest that the Mediator complex is required for sufficient *HNF4A* transcription in both MPHs and AML12 cells.

**Figure 4.**
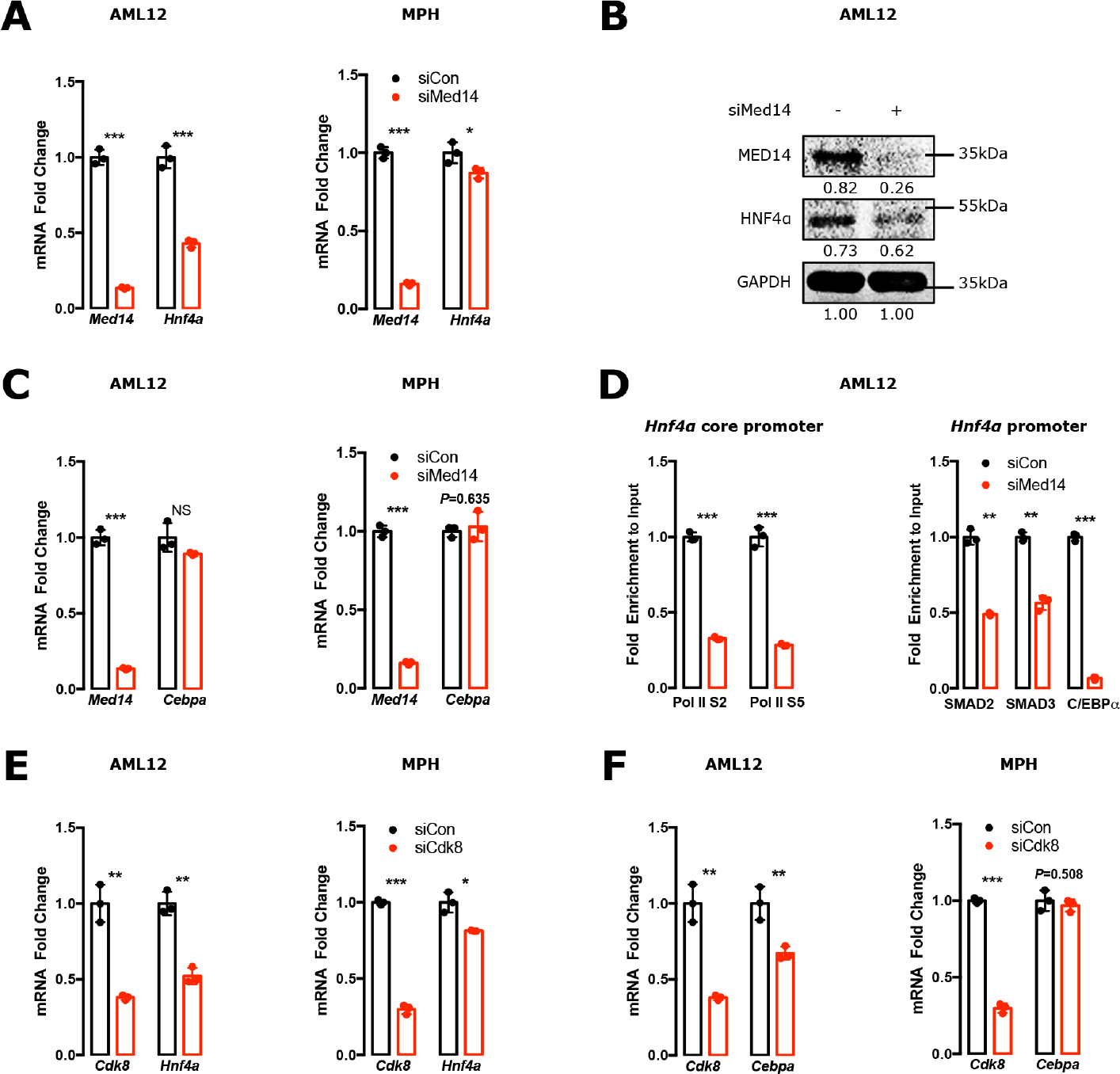
Mediator complex is required for sufficient *HNF4A* transcription. **(A-B)** qPCR (**A**) and western blotting (**B**) examined *Hnf4a* expression in AML12 cells and mouse primary hepatocytes (MPHs). **(C)** qPCR analyzed *Cebpa* expression in AML12 cells and MPHs. **(D)** ChIP-qPCR assays examined the binding activities of Pol II S2 and Pol II S5 on the *Hnf4a* core promoter (-172∼+160bp), and SMAD2, SMAD3, and C/EBPα on the *Hnf4a* promoter (-1750∼-1172bp) in AML12 cells. The cells were transfected by *Med14* siRNA for 48 hours before measurement. **(E-F)** qPCR examined *Hnf4a* (**E**) and *Cebpa* (**F**) expression in AML12 cells and MPHs. The cells were transfected by *Cdk8* siRNA for 48 hours. Bars represent the mean ± SD, *P*-values were calculated by unpaired Student’s t test, *: *P*<0.05; **: *P*<0.01; ***: *P*<0.001; and NS, no significance. Triple experiments were performed, and one representative experiment is shown.

### TGF-*β* reduces *HNF4A* transcription through inhibiting C/EBP*α* expression

SMAD proteins are canonical downstream transcription factors of TGF-*β*^25^. The current finding that SMAD2/3 is required for *HNF4A* transcription seems to conflict with a commonly recognized notion, i.e., that TGF-*β* inhibits HNF4*α* expression in hepatocytes and thus drives EMT^13^. To clarify the relationship between TGF-*β*, SMADs and HNF4*α* in hepatocytes, we initially performed qPCR to examine *Hnf4a* mRNA expression in hepatocytes treated with TGF-*β* for different timepoints. Incubation with TGF-*β* for 2h and 6h significantly induced mRNA expression of *Hnf4a* in MPHs and AML12 cells (**Figure 5A**). However, TGF-*β* treatment for 24h decreased *Hnf4a* mRNA expression (**Figure 5A**). Western blotting showed that TGF-*β*-induced p-SMAD2 was expressed at high levels at all time points measured (**Figure 5B**). In AML12 cells and MPHs, ChIP assays revealed that S5-P Pol II, S2-P Pol II and H3K4me3 binding to the *Hnf4a* core promoter were significantly increased after 2h TGF-*β* incubation, but were remarkably reduced after 24h TGF-*β* administration (**Figure 5C**). Furthermore, TGF-*β* incubation for 2h increased SMAD2, SMAD3 and C/EBP*α* binding to the *Hnf4a* promoter, whereas TGF-*β* incubation for 24h remarkably inhibited C/EBP*α* binding to the *Hnf4a* promoter, although SMAD2 and SMAD3 binding to the promoter was significantly increased at this time point (**Figure 5D**). Ectopic *CEBPA* expression restored TGF-*β*-inhibited HNF4*α* expression in MPHs and HPHs (**Figure 5E-F**). These results suggest that TGF-*β* inhibits *HNF4A* transcription through influencing C/EBP*α* binding to the *HNF4A* promoter.

**Figure 5.**
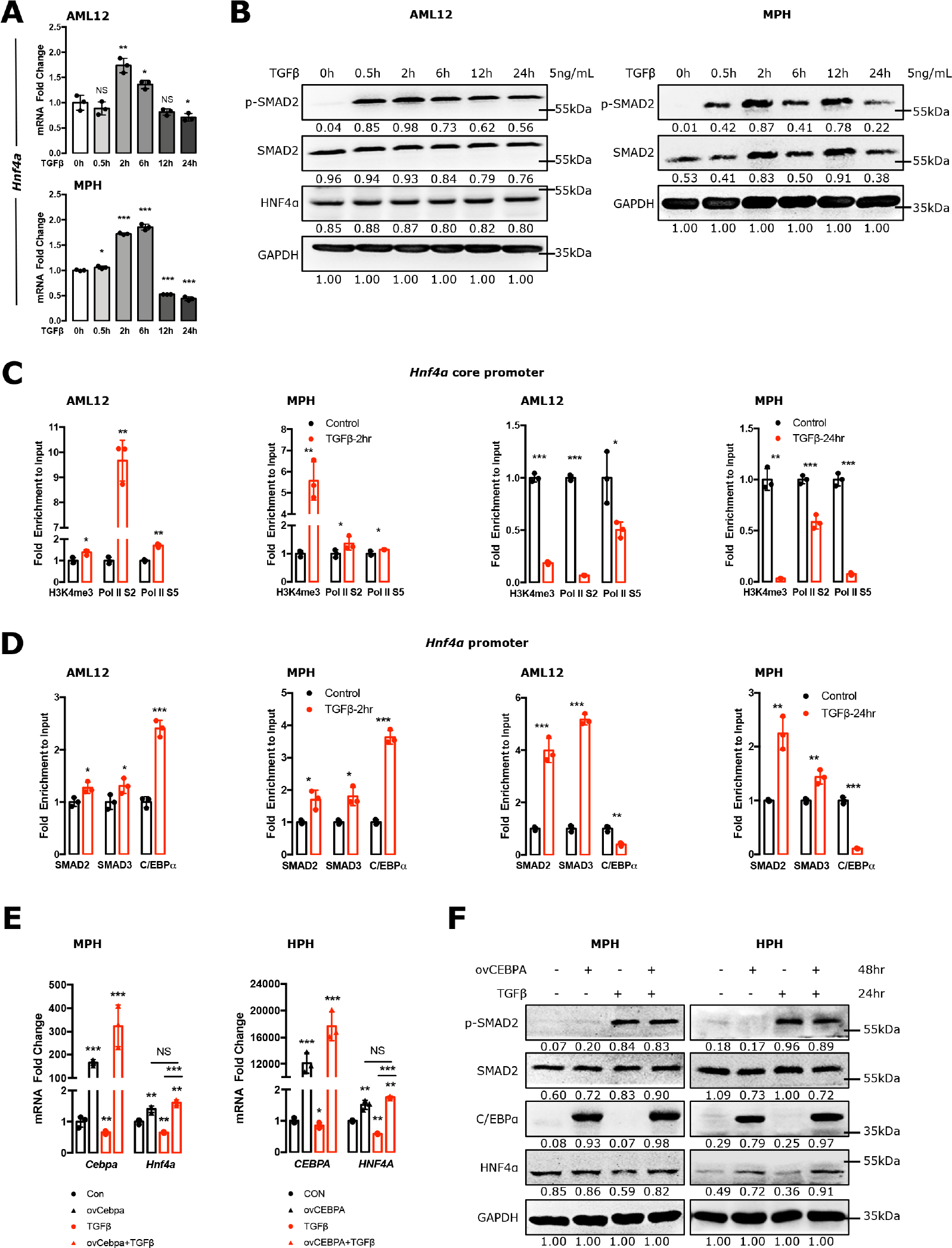
TGF-*β* reduces *HNF4A* transcription through inhibiting C/EBP*α* expression. **(A)** qPCR analyzed a dynamic effect of TGF-*β* (5 ng/mL) on HNF4α expression in AML12 cells and mouse primary hepatocytes (MPHs). **(B)** Western blotting examined p-SMAD2 and HNF4α expression in AML12 cells and MPHs incubated with TGF-*β* for different time points. **(C-D)** ChIP-qPCR assays examined the binding activities of H3K4me3, Pol II S2 and Pol II S5 on the *Hnf4a* core promoter (-172∼+160bp) **(C)**, SMAD2, SMAD3, and C/EBP*α* on the *Hnf4a* promoter (-1750∼-1172bp) **(D)** in AML12 cells and MPHs. The cells were incubated with TGF-*β* for 2 or 24 hours. **(E-F)** qPCR **(E)** and Western blotting **(F)** examined HNF4α expression in MPHs and human primary hepatocytes (HPHs). The cells were transfected by *CEBPA* construct for 48 hours and were subsequently incubated with TGF-*β* for 24 hours. Bars represent the mean ± SD, *P*-values were calculated by One-way ANOVA and unpaired Student’s t test, *, *P*<0.05; **, *P*<0.01, and ***, *P*<0.001. Triple experiments were performed, and one representative result is shown.

### TGF-*β* inhibits C/EBP*α* expression through SMAD binding to the promoter

To clarify how TGF-*β* impacts C/EBP*α* expression, we performed qPCR and Western blotting to examine the effects of TGF-*β* on C/EBP*α* expression in hepatocytes. Although *Cebpa* mRNA expression was temporarily increased following administration of TGF-*β*, C/EBP*α* expression was abolished in MPHs and AML12 cells after 6h of TGF-*β* treatment (**Figure 6A-B**). DNA sequence analysis revealed that the *CEBPA* promoter possesses SMAD2 and SMAD3 binding sites (**Figure 6C**). ChIP sequencing performed in mouse embryonic stem cells and HepG2 cells showed SMAD2 and SMAD3 binding to the *CEBPA* promoter (**Figure 6D**). The binding was also confirmed by ChIP assays in AML12 cells and HPHs (**Figure 6E**). TGF-*β* treatment for 2h reduced SMAD protein binding, but treatment for 24h increased its binding to the *Cebpa* promoter in AML12 cells (**Figure 6F**). Knockdown of *Smad2* or *Smad3* increased mRNA and protein expression of C/EBP*α* in hepatocytes (**Figure 6G- H**). These results suggest that TGF-*β* abolishes *CEBPA* transcription by inhibitory SMAD2/3 binding to the *CEBPA* promoter.

**Figure 6.**
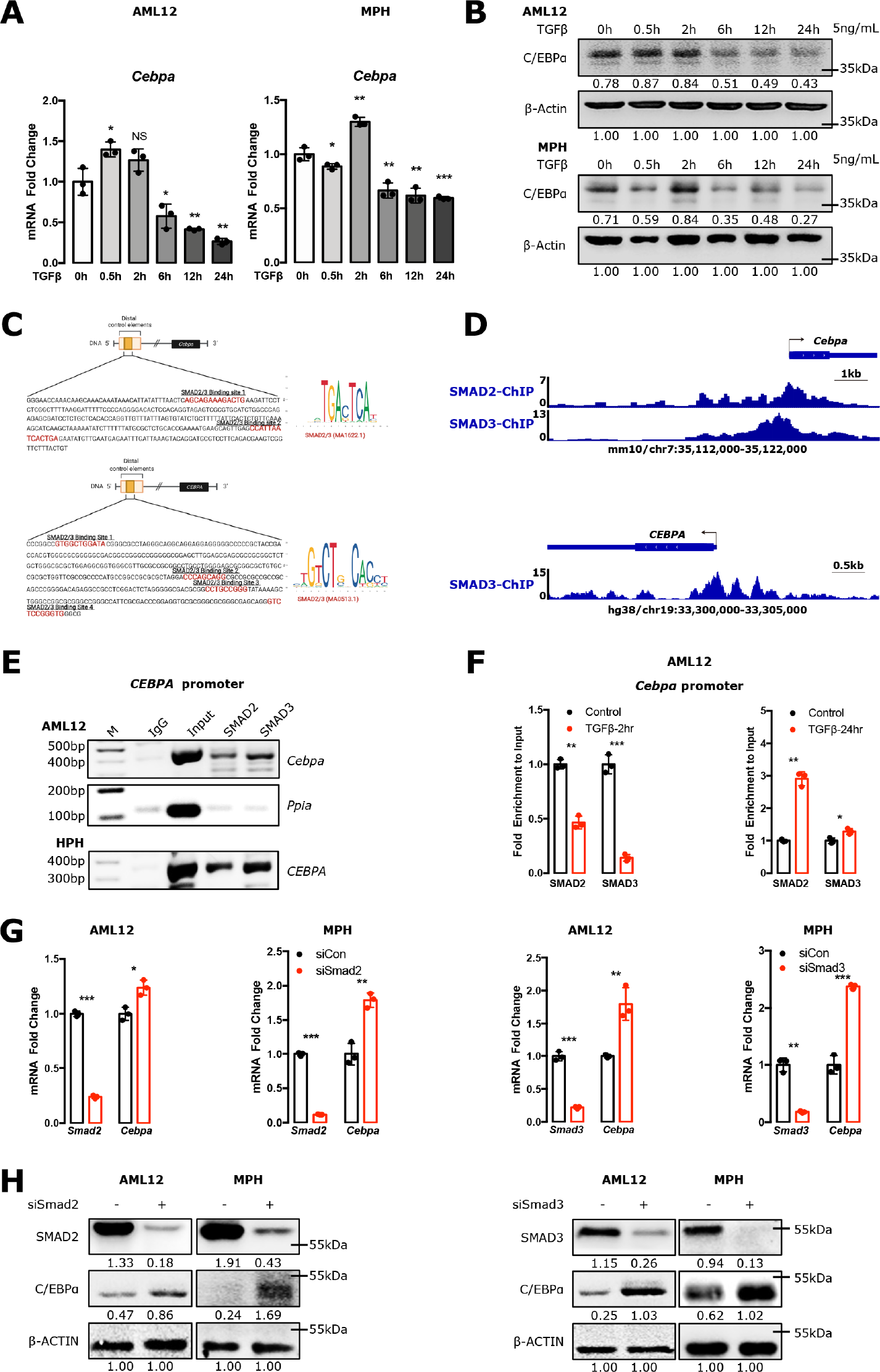
TGF-*β* inhibits C/EBP*α* expression through SMAD2/3 binding to its promoter. **(A-B)** qPCR **(A)** and Western blotting **(B)** analyzed the effect of dynamic TGF-*β* (5 ng/mL) treatment on C/EBPα expression in AML12 cells and mouse primary hepatocytes (MPHs). **(C)** The binding sites of SMAD2/3 on the mouse *Cebpa* (-1997∼- 1509bp) and human *CEBPA* (-1070∼-677bp) promoter was predicted by Jaspar dataset^46^. **(D)** ChIP sequencing showed enrichment profiles of SMAD2 and SMAD3 on *Cebpa* promoter in mouse embryonic stem cells, and SMAD3 on *CEBPA* promoter in HepG2 cells. **(E)** ChIP assays examined SMAD2 and SMAD3 binding to the *CEBPA* promoter in AML12 cells (-1997∼-1509bp) and human primary hepatocytes (HPHs) (- 1070∼-677bp). **(F)** ChIP-qPCR analyzed the binding activities of SMAD2 and SMAD3 on the *Cebpa* promoter (-1997∼-1509bp) in AML12 cells treated with TGF-*β* for 2 or 24 hours. **(G-H)** qPCR **(G)** and Western blotting **(H)** analyzed C/EBPα expression in AML12 cells and MPHs transfected with *Smad2* or *Smad3* siRNA for 24 hours. Bars represent the mean ± SD, *P*-values were calculated by One-way ANOVA and unpaired Student’s t test, *: *P*<0.05; **: *P*<0.01; ***: *P*<0.001; and NS, no significance. Triple experiments were performed, and one representative experiment is shown.

### Insulin is crucial for the maintenance of hepatocellular C/EBP*α* expression *in vivo*

To further clarify the relationship between activated SMAD, C/EBP*α* and HNF4*α* and their roles in damaged livers, we examined their expression levels by immunohistochemical staining (IHC) in liver tissues from 98 patients with chronic HBV infection, cirrhosis, or acute decompensation (AD). Among these patients, hepatic p-SMAD2 levels in 86 patients had been examined previously^16^. In addition, we included a further 12 cirrhotic patients, including 6 with AD. IHC showed HNF4*α* immune positivity in hepatocytes of 74 patients (75.5%) (**Figure 7A**). Analyses based on serial sections further revealed that hepatocytes in 67 (68.4%) of patients simultaneously expressed robust p-SMAD2 and C/EBP*α* (**Figure 7A-B**). In 7 additional patients (7.1%) with HNF4*α* positive reaction, hepatocytes displayed positive p-SMAD2, but undetectable C/EBP*α* expression (**Figure 7A**). There were 22 (22.4%) patients who did not have detectable HNF4*α* levels in hepatocytes (**Figure 7A**). Among them, 5 (5.1%), 15 (15.3%) and 2 (2%) lacked both p-SMAD2 and C/EBP*α*, C/EBP*α*, and p-SMAD2 expression, respectively (**Figure 7A-B**). Impressively, 6 cirrhotic patients with AD did not have detectable levels of any of the three transcription factors (see representative Pat.2 in **Figure 7B**). The remaining 2 patients (2%) without hepatic HNF4*α* expression demonstrated both p-SMAD2 and C/EBP*α* immune reaction in hepatocytes (**Figure 7A**). Taken together, expression of three transcription factors in most examined patients (90.9%) is consistent with the *in vitro* observation: HNF4*α* expression in hepatocytes requires both phosphorylated SMADs and C/EBP*α*.

**Figure 7.**
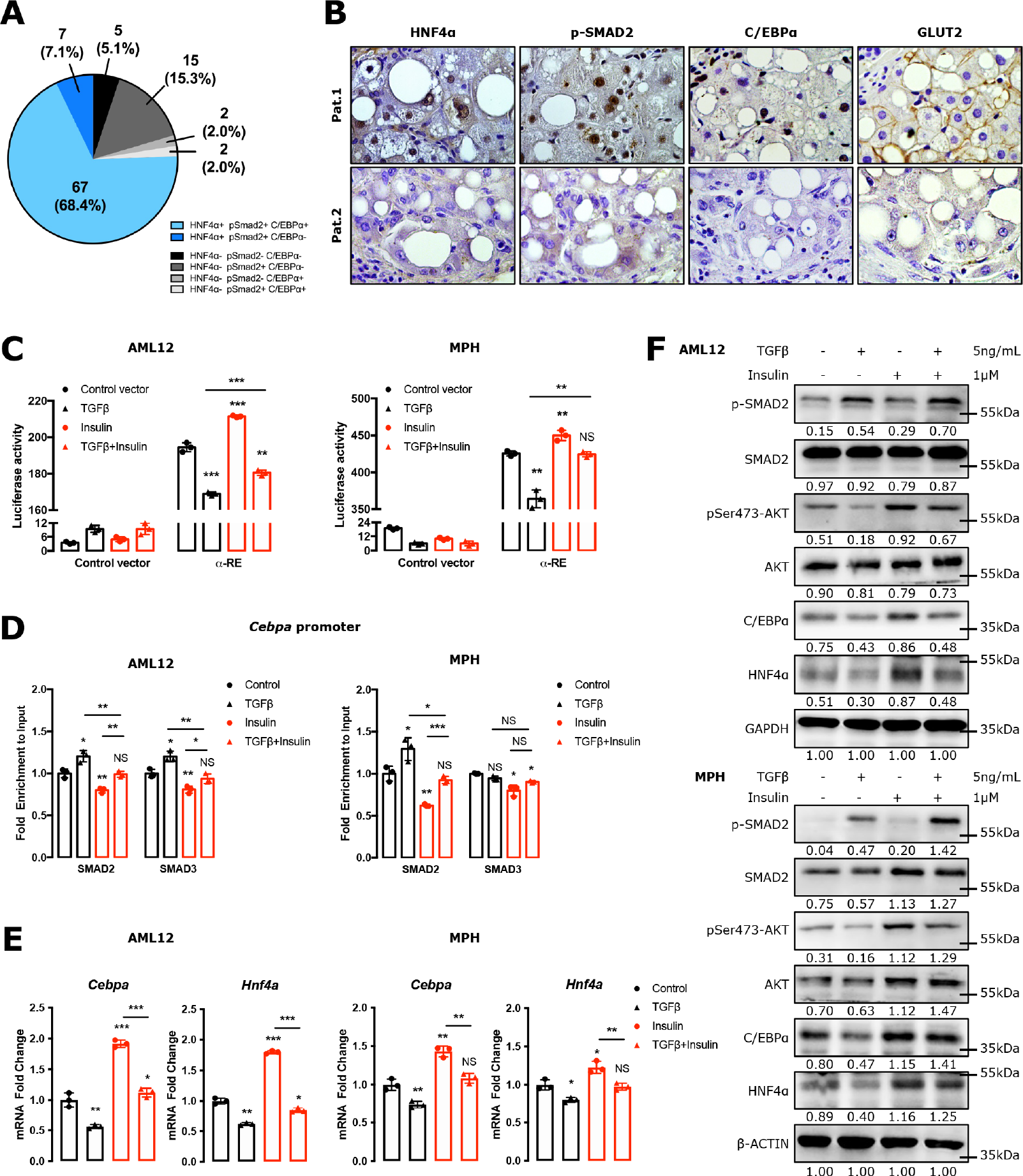
Insulin is crucial for the maintenance of hepatocellular C/EBP*α* expression *in vivo*. **(A)** Immunohistochemical staining for HNF4*α*, p-SMAD2 and C/EBP*α* was performed in 98 patients. Quantification was described in Materials and Methods section. Values are presented as number and percentage. **(B)** HNF4*α*, p-SMAD2, C/EBP*α* and GLUT2 expression is shown in 2 representative cirrhotic patients. Original magnification: x40. **(C)** Luciferase activity of C/EBPα response element (αRE) was measured in AML12 cells and mouse primary hepatocytes (MPHs) transfected with the αRE construct for 48 hours, followed by incubation of TGF-β (5 ng/ml) and/or insulin (1 μM) for 24 hours. **(D)** ChIP-qPCR assays examined the binding activities of SMAD2 and SMAD3 on the *Cebpa* promoter (-1997∼-1509bp) in AML12 cells and MPHs. The cells were treated with TGF-β and/or insulin for 24 hours. **(E-F)** qPCR **(E)** and Western blotting **(F)** analyzed the effect of TGF-β and insulin on C/EBP*α* and HNF4*α* expression in AML12 cells and MPHs treated with TGF-β and/or insulin for 24 hours. Bars represent the mean ± SD, *P*-values were calculated by One-way ANOVA, *: *P*<0.05; **: *P*<0.01; ***: *P*<0.001; and NS, no significance. Triple experiments were performed, and one representative experiment is shown.

The *in vitro* observation that TGF-*β*-induced SMAD activation contributes to HNF4*α* expression, but inhibits C/EBP*α* expression in hepatocytes raises an interesting question: why is hepatic C/EBP*α* expression only lost in patients with AD, but not in patients with chronic HBV infection? We found that insulin prevented TGF-*β*- dependent C/EBP*α* inhibition. Western blotting analyses showed that administration of 1 μM insulin for 30 minutes was sufficient to upregulate protein expression of C/EBP*α* (**Figure S2A**). The upregulated C/EBP*α* levels were maintained for at least 24 hours (**Figure S2A**).

Subsequently, we performed luciferase reporter assay to evaluate the effects of TGF-*β* and insulin on C/EBP*α* response element (*α*RE)^26^. Insulin activated, while TGF-*β* inhibited the activity of *α*RE. Insulin rescued the *α*RE activity from the inhibitory effect of TGF-*β* (**Figure 7C**). ChIP assay shows that insulin inhibited TGF-*β*-dependent SMAD2 and SMAD3 binding to the *Cebpa* promoter (**Figure 7D**). Furthermore, qPCR and Western blotting analyses demonstrated that TGF-*β* was not capable of inhibiting the expression of C/EBP*α* and HNF4*α* when hepatocytes were treated with insulin (**Figure 7E-F**). Inhibition of AKT phosphorylation by pan-AKT inhibitor MK2206 or knocking down *Akt* by RNAi blocked insulin-protected C/EBP*α* and HNF4*α* expression (**Figure S2B-D**). These results suggest a crucial role of insulin in the maintenance of C/EBP*α* and HNF4*α*.

### Insulin resistance leads to loss of C/EBP*α* expression in hepatocytes

These data indicated that systemic inflammatory response-induced hepatic insulin resistance in AD is a major cause of C/EBP*α* and HNF4*α* repression and all its detrimental consequences^27^.

Considering the key role of tumor necrosis factor alpha (TNF-*α*) in the development of insulin resistance, we examined whether TNF-*α*-induced insulin resistance affected C/EBP*α* and HNF4*α* expression in hepatocytes. 1 nM TNF-*α* treatment for 24h inhibited p-AKT expression in hepatocytes (**Figure 8A**). Glucose uptake assay showed that TNF-*α* treatment for 1 day inhibited insulin-dependent glucose uptake in hepatocytes (**Figure 8B**). qPCR assays further revealed that TNF-*α* treatment significantly reduced mRNA expression of solute carrier family 2 member 2 (*Slc2a2*), *Cebpa* and *Hnf4a* in AML12 cells and MPHs (**Figure 8C**), implying TNF-*α*-induced insulin resistance. When hepatocytes were transfected with *Cebpa* construct, TNF-*α*- reduced HNF4*α* expression was partially restored (**Figure 8D-E**).

**Figure 8.**
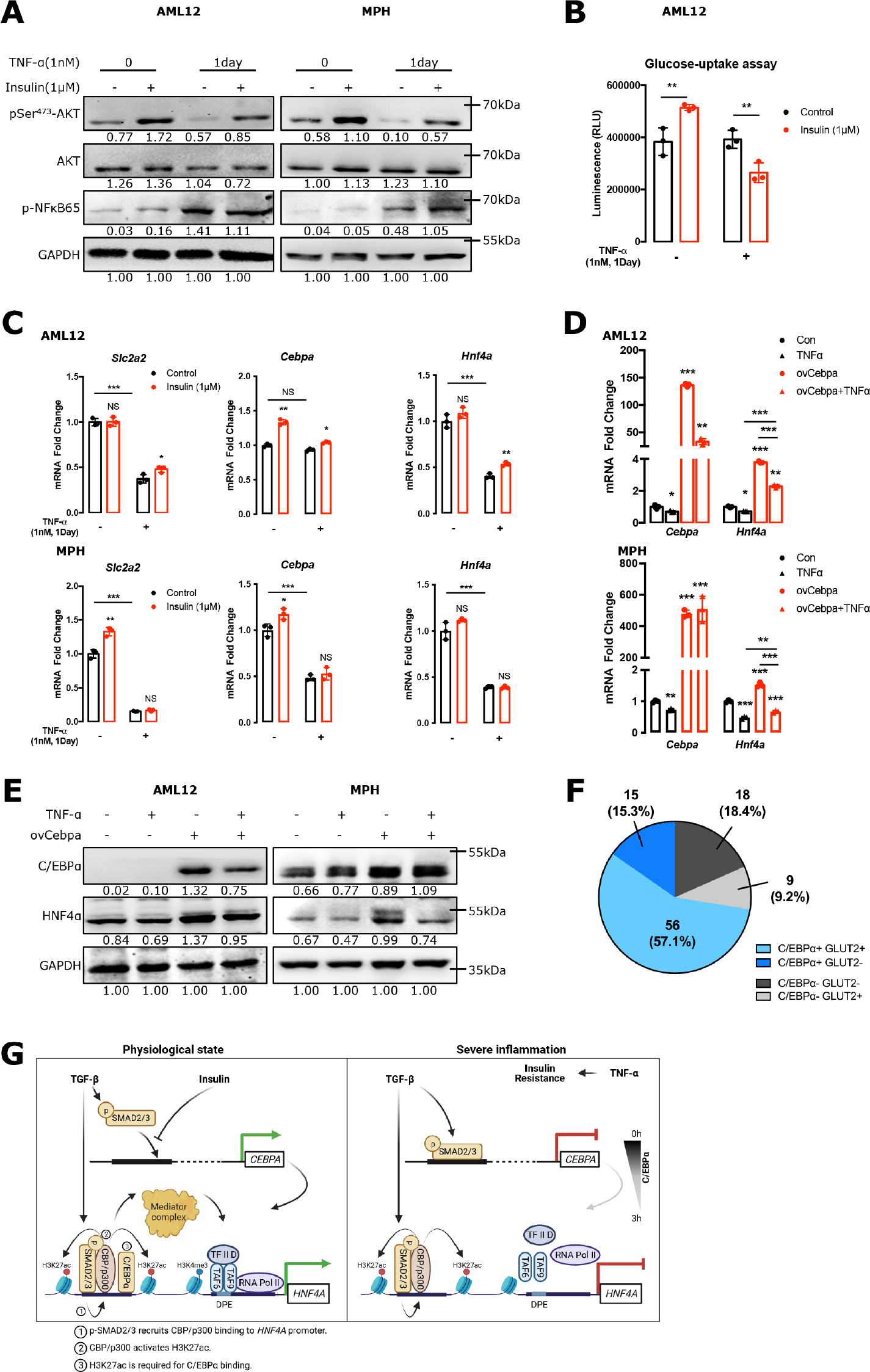
Insulin resistance leads to loss of C/EBP*α* expression in hepatocytes. **(A)** Western blotting analyzed pSer^473^-AKT expression in AML12 cells and mouse primary hepatocytes (MPHs). **(B)** Glucose uptake assay was performed in AML12 cells. The cells were incubated with TNF-*α* (1 nM) for 1 day followed by insulin (1 μM) stimulation for 1 hour. **(C)** qPCR analyzed the effect of TNF-*α* and insulin on *Slc2a2*, *Cebpa*, and *Hnf4a* expression in AML12 cells and MPHs. The cells were incubated with TNF-*α* (1 nM) for 1 day followed by insulin (1 μM) stimulation for 24 hours. **(D-E)** qPCR **(D)** and Western blotting **(E)** analyzed the effect of TNF-*α* and C/EBP*α* on HNF4*α* expression in AML12 and MPHs, the cells were transfected by *Cebpa* construct for 48 hours, followed by incubation with TNF-*α* for 24 hours. **(F)** Immunohistochemical staining for C/EBP*α* and GLUT2 expression was performed in 98 patients. Quantification of immunohistochemical staining was described in Materials and Methods section. Values are presented as number and percentage. **(G)** A schematic diagram summarizes how insulin contributes to the maintenance of C/EBP*α* and HNF4*α* transcription in hepatocytes under TGF-*β* challenge. Bars represent the mean ± SD, *P*-values were calculated by unpaired Student’s t test and One-way ANOVA, *: *P*<0.05; **: *P*<0.01; ***: *P*<0.001; and NS, no significance. Triple experiments were performed, and one representative experiment is shown.

To confirm the impact of insulin signaling on C/EBP*α* expression, we performed IHC for glucose transporter type 2 (GLUT2), the major transporter in charge of hepatocyte glucose uptake^28^, in collected liver tissues. Among 71 patients with hepatic C/EBP*α* expression, 56 had robust GLUT2 expression in hepatocytes while 15 did not (**Figure 7B** and **8F**). In 27 patients without hepatic C/EBP*α* expression, 18 showed undetectable GLUT2, while 9 had GLUT2 expression in hepatocytes (**Figure 7B** and **8F**). In 6 patients with AD, none had detectable C/EBP*α* and GLUT2 immune reaction in hepatocytes (see representative Pat.2 in **Figure 7B**). There were 75.5% of the patients showing positive nuclear C/EBP*α* and membrane GLUT2 expression in hepatocytes simultaneously.

These results strongly suggest a key role of insulin in the maintenance of C/EBP*α* and HNF4*α* expression under TGF-*β* challenge. **Figure 8G** summarizes how insulin contributes to the maintenance of C/EBP*α* and HNF4*α* transcription in hepatocytes under TGF-*β* challenge.

### Insulin prevents hepatocytes from epithelial-to-mesenchymal transition *in vitro*

Given the tight relationship between loss of HNF4*α* and hepatocytes undergoing EMT *in vitro*^11, 14^, the above results led us to ask whether insulin depletion in culture medium results in hepatocytes undergoing EMT. To clarify the role of insulin in EMT, we first measured insulin concentrations in the culture medium of HPHs and AML12 cells at different time points. As shown in **Figure 9A**, insulin concentrations in HPH medium initially were 34.78±1.48 μIU/mL. Over time, insulin concentrations rapidly declined to 12.18±2.66 μIU/mL at 24h, and 0.83±0.49 μIU/mL at 32h, respectively. At 48h, insulin was undetectable in medium (**Figure 9A**). In AML12 culture medium, initial insulin concentrations were surprisingly high (6968.8±751.5 μIU/mL) even though no additional insulin was added. Over time, insulin concentrations rapidly declined to 1585.8±16.7 μIU/mL at 24h, 853.7±6.0 μIU/mL at 48h and 823.1±44.7 μIU/mL at 72h, respectively (**Figure 9A**). Whether hepatocytes were stimulated with TGF-*β* did not alter insulin kinetics in medium (**Figure 9A**).

**Figure 9.**
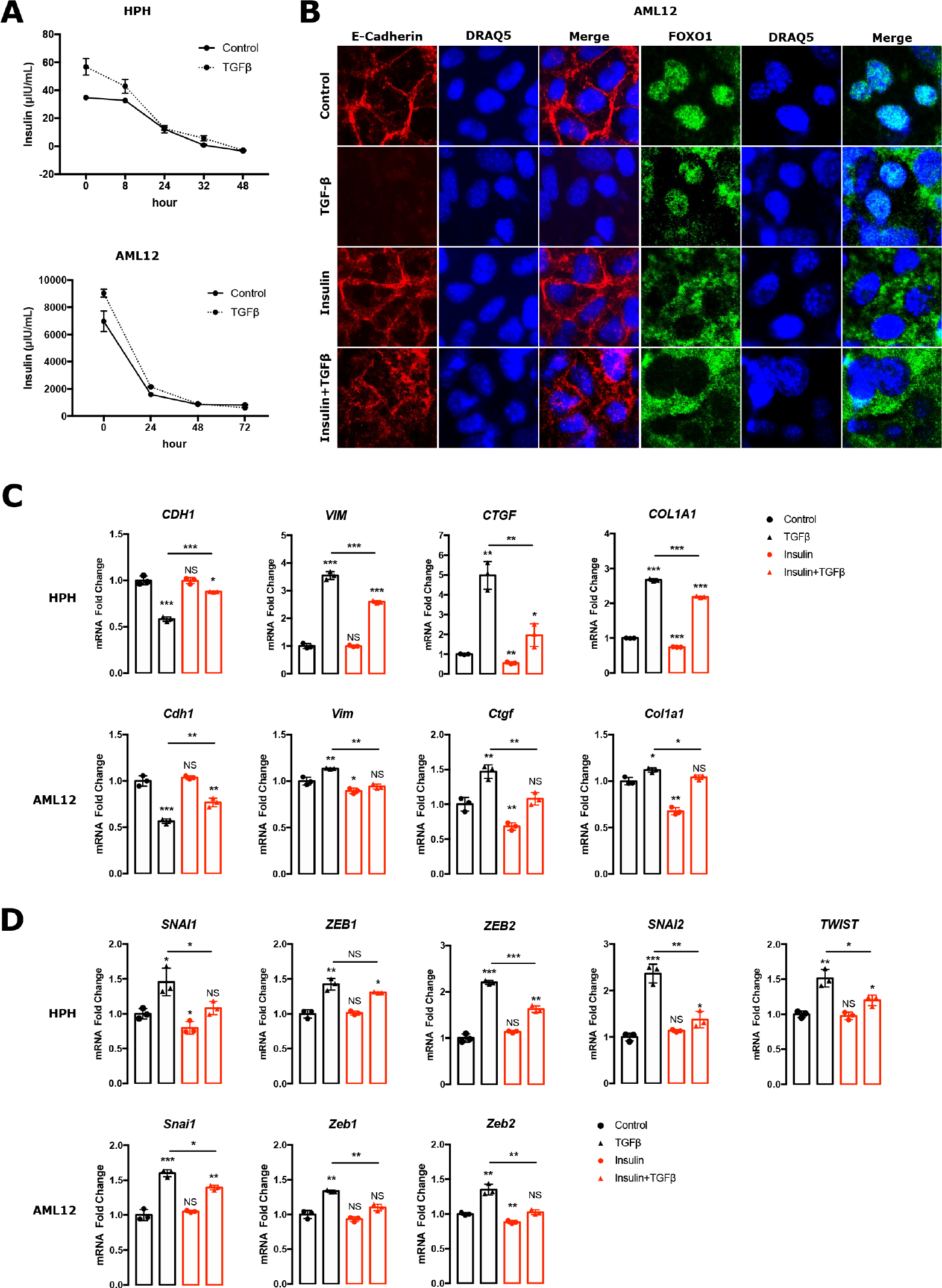
Insulin prevents hepatocytes from epithelial-to-mesenchymal transition *in vitro*. **(A)** Enzyme-linked immunosorbent assay measured insulin concentrations in the culture supernatant of human primary hepatocytes (HPHs) and AML12 at indicated time points. After cell seeding for 24 hours, the cells were changed with fresh medium without insulin. The supernatants were collected at indicated time points. **(B)** Immunofluorescence staining for E-cadherin and FOXO1 was performed in AML12 cells incubated with TGF-β (5 ng/ml) and/or insulin (1μM) for 48 hours. **(C-D)** qPCR was performed to examine expression of epithelial–mesenchymal transition (EMT)- related genes (*CDH1, VIM, CTGF* and *COL1A1*) (**C**) and mesenchymal core transcription factors (*SNAI1/2*, *ZEB1/2*, and *TWIST*) (**D**) in HPHs and AML12 cells incubated with TGF-β (5 ng/ml) and/or insulin (1 μM) for 48 hours. Insulin was added into the culture medium every 2 hours except night (9:00PM – 8:00AM). Bars represent the mean ± SD, *P*-values were calculated by One-way ANOVA and unpaired Student’s t test, *: *P*<0.05; **: *P*<0.01; ***: *P*<0.001; and NS, no significance.

As a commonly recognized *in vitro* model, incubation with TGF-*β* for 48h induces EMT-like phenotype alterations in both collagen monolayer cultured primary hepatocytes or AML12 cells^29^. Therefore, we examined the effect of insulin on TGF-*β*-treated HPHs and AML12 cells. As expected, 5 ng/mL TGF-*β* for 48 hours induced hepatocytes to undergo EMT-like alterations characterized as E-cadherin loss (**Figure 9B**). For comparison, we maintained stable insulin concentration in the medium, by adding 1 μM insulin into the culture medium every 2 hours given that the half-life of insulin is between 20 and 85 minutes^30^. Compared to the hepatocytes with TGF-*β* only treatment, hepatocytes with continuous insulin supplementation maintained E-cadherin expression under TGF-*β* stimulation for 48 hours, and didn’t show induction of TGF-*β*-induced EMT genes, including vimentin (*VIM*), connective tissue growth factor (*CTGF*) and collagen type I alpha 1 chain (*COL1A1*) in HPH and AML12 cells (**Figure 9C**). Immunofluorescence staining further showed that regularly adding 1 μM insulin into the culture medium maintained E-cadherin expression and localization between hepatocytes (**Figure 9B**). Furthermore, insulin administration significantly inhibited TGF-*β*-induced upregulation of core EMT transcription factors such as SNAI1, SNAI2, zinc finger E-box binding homeobox 1 (ZEB1), zinc finger E-box binding homeobox 2 (ZEB2) and twist family bHLH transcription factor (TWIST) in both cells (**Figure 9D**).

### Insulin inhibits TGF-*β*-induced *SNAI1* transcription by impeding nuclear translocation of FOXO1

Next, we scrutinized how insulin controls TGF-*β*-dependent *SNAI1* transcription. ChIP assays revealed SMAD2 and SMAD3 binding to the *SNAI1* promoter in both cells (**Figure 10A**). Binding of SMAD2/3 to the *SNAI1* promoter was inhibited by insulin (**Figure 10B**). Given a low DNA affinity, SMADs usually bind to additional co-factors to form a complex to initiate transcription of target genes^31^. As shown by Gomis et al., Forkhead box O1 (FOXO1) is such a co-factor that is essential for multiple gene activations in cells responding to TGF-*β* challenge^32^. A SMAD-FOXO1 complex was also shown in HaCaT cells^33^. Therefore, we speculated that FOXO1 was also required for SMAD2/3 binding to the *Snai1* promoter in hepatocytes. If so, we hypothesized that insulin might inhibit TGF-*β*-induced SMAD2/3 binding to the *SNAI1* promoter through phosphorylation of FOXO1 and subsequent nuclear exclusion. ChIP assay confirmed that as SMAD2/3, FOXO1 also bound to the neighboring motifs on the *SNAI1* promoter (**Figure 10A**). When we knocked down *FOXO1* by RNAi, TGF-*β*-induced SMAD2 or SMAD3 binding to the *SNAI1* promoter was reduced (**Figure 10C**). TGF-*β*-induced mRNA expression of *Snai1* was remarkably inhibited by *Foxo1* RNAi (**Figure 10D**). It was not surprising that insulin administration induced p-FOXO1 expression and resulted in FOXO1 nuclear exclusion (**Figure 9B, Figure 10E**).

**Figure 10.**
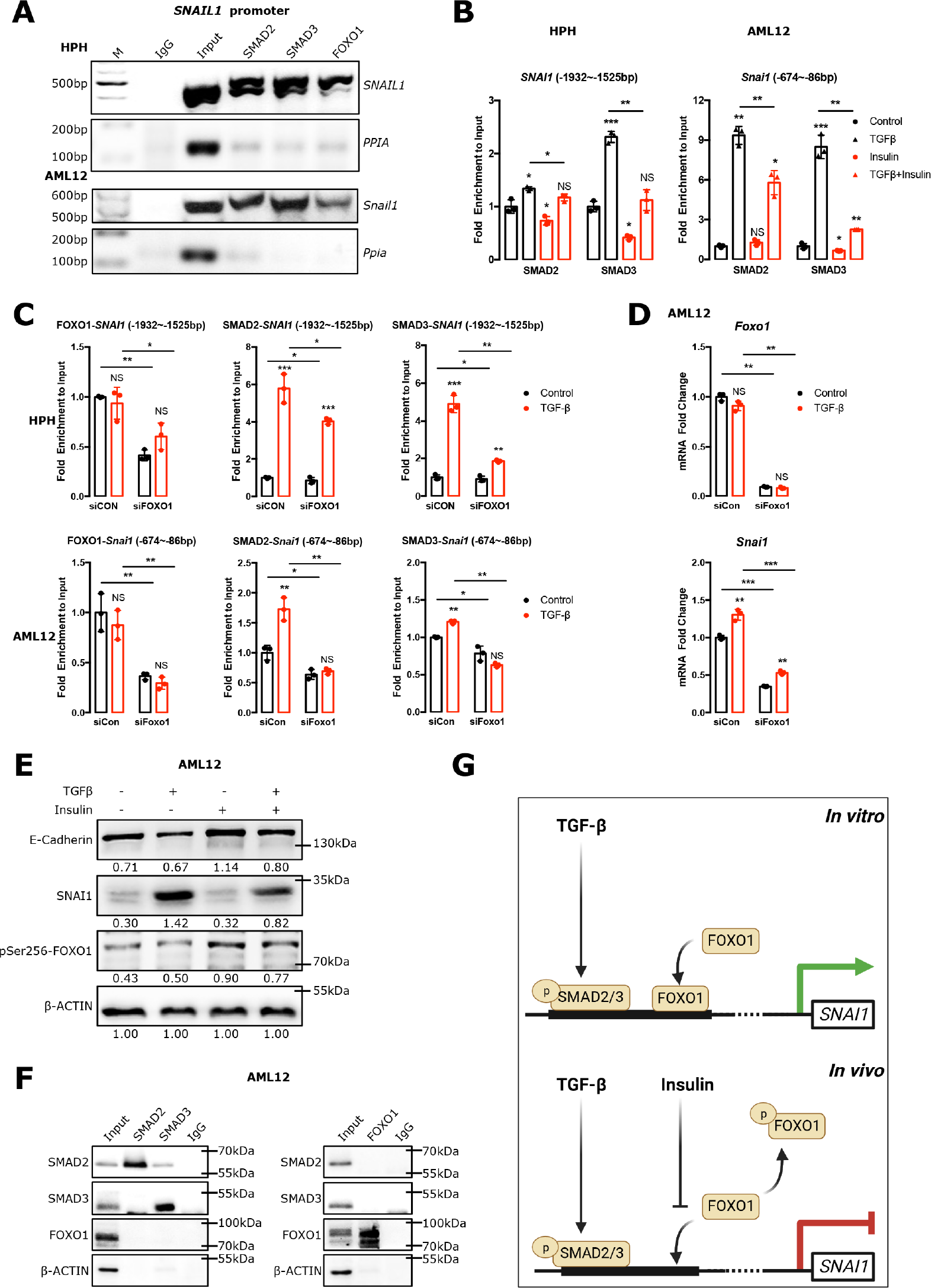
Insulin inhibits TGF-*β*-induced *SNAI1* transcription through impeding unclear translocation of FOXO1 **(A)** ChIP assay showed the binding of SMAD2, SMAD3 and FOXO1 on the *SNAI1* promoter in human primary hepatocytes (HPHs) (-1932∼-1525bp) and AML12 (-674∼-86bp). (**B**) ChIP-qPCR analyzed the binding activities of SMAD2 and SMAD3 on the *SNAI1* promoters in the cells treated with TGF-β (5 ng/ml) and/or insulin (1 μM) for 48 hours. (**C**) The binding activities of FOXO1, SMAD2 and SMAD3 on the *SNAI1* promoter were examined in the cells transfected by siRNA-*FOXO1* for 24 hours, followed by TGF-β (5 ng/ml) stimulation for 24 hours. **(D)** qPCR was performed to measure mRNA expression of *Foxo1* and *Snai1*. **(E)** Western blotting analyzed expression of E-Cadherin, SNAI1 and p-FOXO1 in the cells with indicated treatment. *β*-ACTIN was used as loading control. **(F)** Co-immunoprecipitation examined the protein interactions between SMADs and FOXO1 in AML12 cells. (**G**) A schematic diagram depicts how insulin inhibits TGF-*β*-induced EMT. Insulin was added into the culture medium every 2 hours except night (9:00PM – 8:00AM). Bars represent the mean ± SD, *P*-values were calculated by One-way ANOVA and unpaired Student’s t test, *: *P*<0.05; **: *P*<0.01; ***: *P*<0.001; and NS, no significance.

It is noteworthy that knockdown of *FOXO1* did not significantly alter SMAD2 and SMAD3 binding to the *SNAI1* promoter in hepatocytes without TGF-*β* administration (**Figure 10C**). However, siRNA for *Foxo1* remarkably decreased mRNA and protein expression of SNAI1 in hepatocytes regardless of TGF-*β* treatment (**Figure 10D-E**). Co-immunoprecipitation analysis further showed that SMAD2/3 did not form a complex with FOXO1 in hepatocytes (**Figure 10F**).

These results suggest that both SMAD2/3 and FOXO1 are essential for SNAI1 transcription and expression. However, SMAD2/3 and FOXO1 don’t form a complex to bind to the *SNAI1* promoter in hepatocytes. **Figure 10G** summarizes the process.

## Discussion

Considering the key role of HNF4*α* in performing essential liver functions and the maintenance of hepatocyte identity, we initially focused on how hepatocytes constitutively express HNF4*α* physiologically or under TGF-*β* challenge. We found that the *HNF4A* gene core promoter possesses a DPE box, which is occupied by nucleosomes. Therefore, H3K4me3 is required for providing a harbor for TAF6 and TAF9 that recruit RNA Pol II to initiate its transcription^34^. Sufficient *HNF4A* transcription requires both SMAD2/3 and C/EBP*α* binding to its promoter. SMAD2/3- recruited CBP/p300 activated H3K27ac in the *HNF4A* promoter, which is required for C/EBP*α* binding. A Mediator complex links the active promoter to the RNA Pol II bound core promoter. Disruption of any component of this molecular machinery influences HNF4*α* expression in hepatocytes.

We also address the conundrum that *HNF4A* transcription requires TGF-*β*-activated SMAD2/3, whereas HNF4*α* expression is inhibited by TGF-*β*^13^. These seemingly paradoxical findings are explained by showing that activated SMAD2/3 bind to multiple target genes, including *CEBPA* and *HNF4A*. Although the p-SMAD2/3 complex contributes to *HNF4A* transcription, it equally acts as a transcription repressor of the *CEBPA* gene. As the half-life of C/EBP*α* is between 1-3 hours^35^, C/EBP*α* protein is exhausted when hepatocytes are stimulated with TGF-*β* for extended time periods. Therefore, dynamic observation showed that long-term TGF-*β* incubation inhibits HNF4*α* expression.

TGF-*β*-activated SMAD2/3 inhibits *CEBPA* transcription *in vitro*. However, a large portion of patients simultaneously express robust C/EBP*α*, HNF4*α* and p-SMAD2 in hepatocytes. Why does p-SMAD2 not inhibit C/EBP*α* expression in these patients? We found that insulin inhibits SMAD2/3 binding to the *CEBPA* promoter and thus maintains C/EBP*α* expression under TGF-*β* challenge. In addition, insulin promotes SMAD2/3 binding to the *HNF4A* promoter (data not shown). Patients positive for C/EBP*α* usually show GLUT2 expression in membranes of hepatocytes, indicating insulin sensitivity in these cells, whereas those lacking C/EBP*α* expression often do not have GLUT2 expression. These results suggest that insulin is a key factor protecting C/EBP*α* from TGF-*β* inhibition.

TNF-*α*-induced insulin resistance in skeletal muscle, adipose tissue and hepatocytes is a critical reaction that re-allocates energy from peripheral to central processes in response to severe disease conditions such as sepsis^36^. In this study, we confirmed the impact of TNF-*α* on hepatic insulin handling: Concomitant with insulin resistance, both C/EBP*α* and HNF4*α* expression was inhibited. In patients suffering from ALF and sepsis, hepatocyte nuclei do not show C/EBP*α* and HNF4*α*, while cell membranes do not express GLUT2, indicating hepatic insulin resistance. These results highlight a crucial impact of systemic hormone insulin on hepatocyte function, particularly in ALF and ESLD.

The relationship between insulin, TGF-*b*-SMAD signaling, and HNF4*a* might provide the answer to a longstanding question: why has EMT only been observed in cultured hepatocytes, but not in patients and diseased animals^15^? The answer is that insulin depletion in culture medium causes TGF-*b*-induced EMT *in vitro*. As observed in a substantial number of experiments^13^, TGF-*β* exploits SMAD2/3 to activate core EMT genes such as *SNAI1*. Our study reveals that both SMAD2/3 and FOXO1 binding to the *SNAI1* promoter are required for *SNAI1* transcription. However, in contrast to HaCaT cells^33^, SMAD2/3 do not interact with FOXO1 in hepatocytes. In this cell type insulin induces nuclear exclusion of FOXO1, and thus reduces SNAI1 expression. In cultured HPHs, insulin concentrations in medium sharply decline from around 35 μIU/mL to undetectable levels within 48h. In AML12 cells, the culture medium contains extremely high insulin levels (around 7000 μIU/mL). As in HPHs, the concentrations rapidly decline to around 800 μIU/mL within 48h. We define such an insulin reduction as “insulin depletion”. Following insulin depletion in culture medium, TGF-*β*-SMAD signalling initiates the transcription of core EMT genes and thus causes an EMT-like phenotype of hepatocytes. Regularly adding insulin to the culture medium completely inhibits this phenotype alteration. As a master systemic regulator of energy metabolism, insulin concentrations are tightly controlled in mammals^37^. Insulin depletion is a state that must never occur in both humans or animals, as it would result in rapid death. This might explain why complete hepatocyte EMT has never been observed *in vivo*.

To date, the crosstalk between TGF-*β* and PI3K pathway in different biological processes including EMT has been extensively investigated. Coordination between TGF-*β* and insulin-PI3K-AKT pathways are essential for EMT in cancer cells (37–40). However, we show an opposite effect between TGF-*β* and insulin in hepatocytes. As an essential signal regulating anabolic metabolism, the response to insulin stimulation is determined by the cell type as well as the actual physiological or disease condition. In mammals, only hepatocytes, adipose cells, and skeletal muscle cells develop insulin resistance during sepsis to ensure sufficient energy supply to priority organs. Therefore, it is not surprising that hepatocytes demonstrate a unique response to insulin and TGF- β stimulation. The special cell context of hepatocytes should be further investigated to clarify whether inhibition of TGF-β-induced EMT by insulin is specific for this cell type.

## Materials and methods

Chemical reagents, primers and antibodies used in this study are summarized in Supplementary Table 1-5.

### Patients

Samples were collected from 98 patients, comprising 56 non-cirrhotic and 42 cirrhotic patients at the Department of Medicine II, University Medical Center Mannheim, Medical Faculty Mannheim, Heidelberg University, and the Department of Gastroenterology and Hepatology, Beijing You’an Hospital, Affiliated with Capital Medical University. Among 42 cirrhotic patients, 30 were compensated cirrhosis while 12 suffered from acute decompensation (AD) and received liver transplantation.

The study protocol was approved by local Ethics Committees (Jing-2015-084, and 2017-584N-MA). Written informed consent was obtained from patients or their representatives. A portion of patients have been investigated in a previous study^16, 38^.

Patients with AD were defined by the development of one or more major complications of liver diseases: (i) development of grade 2 to 3 ascites within <2 weeks; (ii) hepatic encephalopathy; (iii) gastrointestinal hemorrhage; (iv) bacterial infections (spontaneous bacterial peritonitis, spontaneous bacteremia, urinary tract infection, pneumonia, cellulitis)^39^.

### Cells

Mouse primary hepatocytes were isolated by collagenase (C2-28, Merck) digestion of livers from *ad libitum* fed mice using two-step collagenase perfusion technique as described previously^40, 41^. The cells were seeded on collagen-coated plates in Williams’ E medium supplemented with 10% Fetal bovine serum (FBS), 2mM L-glutamine, penicillin (100 U/mL)-streptomycin (100 μg/mL), 100 nM dexamethasone, and 5% of Insulin-Transferrin-Selenium (ITS). Cells were cultured in a 37°C incubator with 5% CO2 overnight before treatment. Primary human hepatocytes were isolated by the Cell Isolation Core Facility of the Biobank Großhadern, University Hospital, LMU Munich. These cells were seeded in adhesion medium containing Williams’ E medium supplemented with 10% FBS, 100 nM dexamethasone, 10 mg/ml insulin, 2mM L-glutamine and penicillin (100 U/mL)-streptomycin (100 μg/mL) in a 37°C incubator with 5% CO2.

AML12 cells were kindly provided by Dr. Nelson Fausto. Cells were grown in DMEM/F-12 medium (21331-020, Gibco) supplemented with 10% FBS, 5% ITS, 2 mM L-glutamine, penicillin (100 U/mL)-streptomycin (100 μg/mL), 100 nM dexamethasone at 37°C in a humidified atmosphere containing 5% CO2.

Before TGF-β treatment, hepatocytes were serum starved for 8 hours. For experiments relevant to insulin incubation, no additional insulin was added into the medium.

HEK293 cells were used to perform luciferase reporter assay. Cells were cultured in DMEM medium (BE12-709F/12-M, Biozym) supplemented with 10% FBS, 2 mM L-glutamine and penicillin (100 U/mL)-streptomycin (100 μg/mL). Cells were grown at 37°C in a humidified atmosphere containing 5% CO2.

### Immunohistochemistry

Liver tissue sections were dewaxed in xylene (9713.2, Carl Roth) and rehydrated in an ethanol gradient (K928.4, Carl Roth). For antigen retrieval, slides incubated with 1 mM Ethylene Diamine Tetraacetic Acid solution (EDTA, E9884, Sigma-Aldrich, pH 8.4) or citrate acid buffer (C2404, Sigma-Aldrich, pH 6.0) were heated by microwave (1000 W, 15 seconds full power, 45 seconds shut down, 10 cycles). Following cooling at room temperature (RT) for 30 mins, the slides were incubated in DAKO blocking peroxide (S200389-2, DAKO) for 30 mins to eliminate non-specific staining. Subsequently, primary antibodies were incubated overnight at 4°C. Next day, following 3 washes with phosphate-buffered saline (PBS, L182-50, Biochrom), sections were incubated with the horseradish peroxidase-labelled secondary antibodies for 45 mins. Visualization was achieved with 3,3-diaminobenzidine (DAB, D5905, Sigma-Aldrich). Counterstaining was performed with hematoxylin (H-3401-500, Vector Laboratories). Finally, all sections were dehydrated, mounted with mounting medium (C9368, Sigma-Aldrich). Images were taken under a microscope (DMRE, Leica).

### Immunofluorescence

Cells were fixed in ice-cold 4% paraformaldehyde for 15 mins, followed by permeating with 0.4% Triton X-100 in PBS for 5 mins, and blocking with 0.025% Triton X-100 and 1% BSA for 1 hour at room temperature with shaker. Subsequently, the cells were incubated with primary antibodies overnight at 4 °C. Next day, the cells were probed with fluorochrome-conjugated secondary antibody for 1 hour at room temperature followed by washing with PBS three times. Nuclei were identified with DRAQ5 prepared in 1%BSA for 10 mins. The sections were photographed with a Leica Confocal fluorescence microscope.

### Chromatin immunoprecipitation and quantitative real-time PCR

Chromatin immunoprecipitation and quantitative real-time PCR (ChIP-qPCR) analyses were performed as described previously with minor modifications^42^. Briefly, for each ChIP, cells were isolated from one 10 cm cell culture dish (95% confluence). Cells were washed twice with PBS (14190144, Thermo Fisher) and incubated with 1% formaldehyde at 37 °C for 10 min to crosslink proteins and DNA. Following ice-cold PBS washing twice, the cells were collected in lysis buffer (1% SDS (2326.2, Carl Roth), 5 mM EDTA in 50 mM Tris, pH 8.1) containing 1% protease inhibitor cocktail. The chromatin was sonicated by 30 seconds pulses, followed by 30 seconds interval to produce DNA fragments with an average of 200-500 bp length. After sonication, the samples were centrifuged at 14000 rpm, 4°C for 10 mins. The supernatants were collected in a clean tube and diluted by adding dilution buffer. 5μg of the antibody or normal IgG and 50μL of agarose beads were added into the samples. After rotating overnight at 4°C, the beads were collected by centrifugation at 2000 rpm for 10 mins. Then the pellets were successively washed with 1 ml TSEI (0.1% SDS, 1% Triton X- 100, 2 mM EDTA, 20 mM Tris-HCl Ph8.0, 150 mM NaCl), TSEII (0.1% SDS, 1%

Triton X-100, 2 mM EDTA, 20 mM Tris-HCl Ph8.0, 500 mM NaCl), BufferIII (0.25 M LiCl, 1% NP-40, 1% Deoxycholic acid, 10 mM Tris-HCl Ph8.0) and TE buffer (10 mM Tris-HCl Ph8.0, 1 mM EDTA) for 10 minutes each at 4°C. After centrifugation at 2000 rpm for 10 minutes, the beads were resuspended in elution buffer and kept washing at room temperature for 10 mins. Immunoprecipitated complexes were de-crosslinked by heating to 65 °C for 6h. The pulled-down DNA fragments were extracted and purified using phenol-chloroform extraction/ethanol precipitation. The samples were subjected to quantitative PCR using SYBR green assay (A25918, Thermo Fisher). The primer sets used for these assays are listed in **Table S1**.

### RNA extraction and quantitative real-time PCR

All reagents and containers used for RNA processing were RNase-free grade or treated with 0.1% DEPC (4387937, Thermo Fisher) to eliminate RNase contaminants. Total RNA was extracted from cells and tissues in TRIzol (15596018, Invitrogen) according to the manufacturer’s instructions. Reverse transcription of 500 ng RNA to synthesize cDNA was performed using random primers (SO142, Thermo Fisher) and RevertAid H Minus Reverse Transcriptase (EP0452, Thermo Fisher) according to the manufacturer’s instructions. The quantitative real-time PCR (qPCR) assays were carried out on a StepOnePlus system (Applied Biosystem) using SYBR Green Master Kit (A25918, Thermo Fisher). Primers for qPCR were listed in **Table S1**. Three biological replicates per condition were measured. The relative abundance (fold changes) of each target gene compared with a set of internal controls were determined by the -2^ΔΔCT^ method^43^.

### Western Blotting

Cells were lysed in RIPA buffer (R0278, Sigma-Aldrich) supplemented with protease inhibitors (36978, Thermo Fisher) and phosphatase inhibitors (P5726, Sigma-Aldrich) on ice for 10 mins. Cell lysates were collected by centrifugation at 14000 rpm, 4°C for 15 mins. The total protein concentration in the supernatant was determined by a Bio-Rad protein assay kit (5000113-115, Bio Rad, Bradford method). Following the addition of LDS sample buffer (4x, NP0007, Lifetechnologies), the samples were boiled at 90°C for 10 mins. Twenty microgram proteins were separated by 10-15% SDS-PAGE and transferred onto 0.2 μm nitrocellulose membranes (4685.1, Carl Roth). Following blocking with 5% non-fat milk for 1 hour at room temperature, the membranes were incubated with primary antibodies at 4°C overnight. Next day, the membranes were washed with tris buffered saline plus 0.05% Tween 20 (TBST) and incubated with secondary antibodies for 1 hour. After washing with TBST for three times, the membranes were incubated and visualized by the Pierce™ ECL Plus Western Blotting Substrate (32132, Lifetechnologies).

### siRNA transfection

Confluent hepatocytes (80% MPHs or 60% AML12) were seeded 24 hours before transfection. siRNAs were transfected into cells using Lipofectamine RNAiMAX reagent (13778-075, Thermo Fisher) according to the manufacturer’s instruction. The transfected cells were incubated at 37 °C for 24 to 48 hours, followed by extraction of cellular DNA, RNA and proteins. A non-targeting negative stealth siRNA (scrambled, sc-37007, Santa Cruz) was used as a negative control. The sequences of siRNA are listed in **Table S4**.

### Plasmid transfection

Cells were seeded in 12-well plates and grown to 90∼95% confluency. The transfection was performed using Lipofectamine 3000 reagent (L3000008, Thermo Fisher) following the manufacturer’s instruction. 1.75 μg/well of plasmids were transfected into cells. Empty plasmid p-Flag-CMV4, pScalps were used as a control vector. After the cells were incubated at 37 °C for 48 hours, cellular RNA, DNA and proteins were extracted.

### Luciferase reporter assay

Hepatocytes were cultured in standard medium without insulin. C/EBPα response element (αRE; CGCGTATTGGCCAATATTGG CCAATCTCGA) was transfected into hepatocytes using lipofectamine 3000 reagent (L3000008, Thermo Fisher). The αRE construct was described previously^26^. Cells were treated with insulin (1 μM), TGF- β (5 ng/mL), or both for 24 hours. The luciferase assay was performed according to the manufacturer’s instruction (E1500, Promega).

Mouse genetic *Hnf4a* promoter (-2000 to 0) and its deletion mutants (-350 to 0) were cloned into pGL4.14 vector (E6691, Promega) by Gibson Assembly kit (A46624, Thermo Fisher). Primer sequences are listed in **Table S1**. In brief, HEK293 cells were transfected with pGL4.14-*Hnf4a* plasmids or pGL4.14-vector plasmids as the control plasmid with lipofectamine 3000 reagents. For luciferase reporter assay, the pGL4.74 plasmid (E6921, Promega) was used as a loading control to standardize the dual-luciferase activity. Then cells were co-transfected with pGL4.14-*Hnf4a* plasmids and *Smad2*/*Samd3*/*Cebpa* plasmid for 48 hours. Dual-Luciferase Assay Kit (E2920, Promega) was used to measure the dual-luciferase activity according to the manufacturer’s instruction.

### Glucose uptake assay

Cells were seeded in a 24-well plate and were cultured in the medium without insulin. Twenty-four hours later, the cells were incubated with TNF-*α* (1 nM) for 1 day at 37°C and 5% CO2. Then insulin (1 μM) was added into the medium for 1 hour. After removing the supernatant and washing with PBS, the cells were treated by 2- deoxyglucose (2-DG) for 10 mins at room temperature. The cells were successively lysed and assayed by a 2-DG uptake assay kit (J1341, Promega) according to the instruction. Total 2-deoxyglucose-6-phosphate (2DG6P) was measured by recording luminescence using a 0.3-1 sec integration on a luminometer.

### Enzyme-linked immunosorbent assay

The insulin enzyme-linked immunosorbent assay (ELISA) kits were purchased from Thermofisher (EMINS) and Sigma (RAB0327). 100 μl standard solution or samples were added into each well. The plates were incubated for 2 hours at RT with continuous shaking. Followed by washing with 200 μl of wash buffer five times, 100 μl of biotinylated albumin antibody were added and incubated for 1 hour. After washing, the plates were added with 100 μl of streptavidin-peroxidase conjugate and incubated for 45 mins. Followed by additional washing, 100 μl of chromogen substrate were added and incubated for 30 mins. When the color density developed, 50 μl of stop solution were added. Plates were immediately read at 450 nm wavelength by a microplate reader. A standard curve based on the serial dilutions of data with concentrations on the X axis (log scale) *versus* absorbance on the Y axis (linear) was generated. The concentrations were calculated according to the standard curve.

### Immunoprecipitation analysis

Cells were harvested with ice-cold PBS and lysed in Lysis Buffer (50 mM Tris-HCl [pH 7.4], 0.1% NP-40, 100 mM NaCl, and 1 mM EDTA) containing protease inhibitors (S8820, Sigma-Aldrich) and phosphatase inhibitors (P5726, Sigma-Aldrich). The cell extracts were pre-cleaned with protein A/G magnetic beads (88804, Thermo Fisher Scientific) at 4℃ for 60 min. The samples were then incubated with anti-SMAD2, anti- SMAD3, anti-C/EBPα, and anti-FOXO1 antibodies or normal rabbit IgG at 4℃ overnight and subjected to SDS-PAGE by western blot analysis.

### Data Availability

ChIP-seq data of mESC and HepG2 cells were obtained from the GEO (accession no. GSE 125116, GSE 169790)^44, 45^.

### Statistical analysis

Analyses were performed using SPSS Statistics 23.0. The unpaired Student’s t test was used to determine statistical significances between two groups. One-way ANOVA was used to test the statistical differences among multiple comparisons. *P* values less than 0.05 were considered significant and represented graphically as *, *P*<0.05; **, *P*<0.01; ***, *P*<0.001, unless otherwise indicated.

## Acknowledgements

We are grateful to Dr. Nelson Fausto for kindly providing AML12 cells and Dr. Atsushi Miyajima for *α*-RE construct. We thank Drs. Atsushi Miyajima and Peter ten Dijke for constructive discussion. We also thank the Human Tissue and Cell Research Foundation, a nonprofit foundation regulated by German civil law, which facilitates research with human tissue through the provision of an ethical and legal framework for prospective sample collection. We acknowledge the support of the LIMA Live Cell Imaging at Microscopy Core Facility Platform Mannheim (CFPM).

## Author’s contributions

H.L.W. conceived and designed the project. R.F., S.M., N.H., H.L., C.S., and H.D. collected the patient samples and performed pathological experiments. R.F., C.T., T.L., Y.L., C.S., K.K., S.S.W., S.W., X.F.L., R.L., and S.W. undertook the *in vitro* experiments. R.F. and H.L.W. drafted the article. R.F., C.M., R.L., M.P.A.E., S.D., H.D., H.W., and H.L.W. discussed the data and edited the article critically.

