## Supplemental tables and figures for "Insulin determines the effects of TGF-β on HNF4α transcription and epithelial-to-mesenchymal transition in hepatocytes"

**Supplementary tables**

**Table S1. Primers used in the study**

**ChIP-Primers**

| Gene | Forward primers (5'-3') | Reverse primers (5'-3') |
|---|---|---|
| *HNF4A* (-250~+67bp) | TTGGGGTTGGCTCTCTAGGA | CAGGCAGATGTGGAGTCAGG |
| *Hnf4a* (-172~+160bp) | CCTGGTTCCAAGAAGCCACT | ATCTGCCAGGTGATGCTCTG |
| *Hnf4a* (-350~0bp) | CCACTCTCCCTCCCTCCTCT | TTATCTTATTGATTCTTCTAAT CACCCAAGGTGG |
| *Hnf4a* (-214~-58bp) | CCTCTCTCAAGTTGGAAGGCT | TCACTTAGGGAACCTGTGGC |
| *Hnf4a* (-1750~-1172bp) | TGGCTCTCTCTGTCCTCTCC | AAAATGGGGTGACGATGGCT |
| *CEBPA* (-1070~-677bp) | ACGCAGGCAGGTAAAGCTAA | TCAAGAACCCACCCAGCATC |
| *Cebpa* (-1997~-1509bp) | GTGTTCCCTTTCCCCTCAGG | TCTAGGGTCGCAGGTCAAGA |
| *SNAI1* (-1932~-1525bp) | ATGAAAGGAAGCCAGCGTGA | TCGTCCTCCCCTGGTTCTAG |
| *Snai1* (-674~-86bp) | TTTCCCCTCGGTGCTTCTTC | ACTACGATCCCCTAGCAGCA |
| *PPIA* | AGCATGTGGTGTTTGGCAAA | TCGAGTTGTCCACAGTCAGC |
| *Ppia* | GAGCTGTTTGCAGACAAAGTT | CCCTGGCACATGAATCCTGG |

**qPCR-Primers**

| Gene | Forward primers (5'-3') | Reverse primers (5'-3') |
|---|---|---|
| *Ppia* | AGGATTCATGTGCCAGGGTG | GCCATCCAGCCATTCAGTCT |
| *Hnf4a* | AGATGCTTCTCGGAGGGTCT | GCCACTCACACATCTGTCCA |
| *Cebpa* | GCAAAGCCAAGAAGTCGGTG | CACCTTCTGTTGCGTCTCCA |
| *Smad2* | TCCGGCTGAACTGTCTCCTA | CTGTGACGCATGGAAGGTCT |

| | | |
|---|---|---|
| *Smad3* | AGAGGTGTGCGGCTCTACTA | GGGCAGCAAATTCCTGGTTG |
| *Taf6* | CACTCACCATCACACAGCCT | AGGAGGAGAAGGCTGAGGAG |
| *Taf9* | ACCCCTTTGCCACTGATCAA | TGGAGTCGGTGTACCTAGGG |
| *Cdk8* | CCTCCGACTATCAGCGTTCC | GCTGAGTATCCCATGCTGCT |
| *Med14* | AGGGGCCAGTTTTGCTAGTC | CAGTTTCCTGCTCGTCCACT |
| *Slc2a2* | GCATCGACTGAGCAGAAGGT | CTCCACAAGCAGCACAGAGA |
| *Cdh1* | CAGGTCTCCTCATGGCTTTGC | CTTCCGAAAAGAAGGCTGTCC |
| *Vim* | GCGCCCTCATTCCCTTGTTGCA | GGCGCAGGGCATCGTTGTTC |
| *Ctgf* | TGTGTGACGAGCCCAAGGA | TTGGGTCTGGGCCAAATGT |
| *Col1a1* | GGAGAGAGCATGACCGATGG | AAGTTCCGGTGTGACTCGTG |
| *Snai1* | GTGAGAAGCCATTCTCCTGCT | CTCTTCACATCCGAGTGGGTT |
| *Zeb1* | CCAGAAGCCAGCAGTCATGA | GGAGCAGCTGAAGTTGTCCT |
| *Zeb2* | CAACCATGAGTCCTCCCCAC | TTCTGGCCCCATTGCATCAT |
| *Foxo1* | ATCTGCCATGAACCGCTTGA | TCACAGTCCAAGCGCTCAAT |
| *GAPDH* | GGAGCGAGATCCCTCCAAAAT | GGCTGTTGTCATACTTCTCATGG |
| *HNF4A* | CACGGGCAAACACTACGGT | TTGACCTTCGAGTGCTGATCC |
| *CEBPA* | TATAGGCTGGGCTTCCCCTT | AGCTTTCTGGTGTGACTCGG |
| *SMAD2* | CCAGGAATTTGCTGCTCTTC | TCCATAGGGACCACACACAA |
| *SMAD3* | GGCTCCCTCATGTCATCTACT | AGTAGGTAACTGGCTGCAGGT |
| *CDH1* | CGAGAGCTACACGTTCACGG | GGGTGTCGAGGGAAAAATAGG |
| *VIM* | TGGCCGACGCCATCAACACC | TGGCGCAGGGCGTCATTGTT |
| *CTGF* | TTGGCCCAGACCCAACTA | GCAGGAGGCGTTGTCATT |

| | | |
|---|---|---|
| *COL1A1* | CGGTGTGACTCGTGCAGC | ACAGCCGCTTCACCTACAGC |
| *SNAI1* | AATCGGAAGCCTAACTACAGCG | GTCCCAGATGAGCATTGGCA |
| *SNAI2* | GCTGGCCAAACATAAGCAGC | AGGGTCTGGAAAACGCCTTG |
| *TWIST* | TGCATGCATTCTCAAGAGGT | CTATGGTTTTGCAGGCCAGT |
| *ZEB1* | GCCAACAGACCAGACAGTGT | CCCCAGGATTTCTTGCCCTT |
| *ZEB2* | GGAGACGAGTCCAGCTAGTGT | CCACTCCACCCTCCCTTATTTC |

**Luciferase-Primers**

| **pGL4.14** | Primer sequence (5'-3') |
|---|---|
| *Hnf4a* P-2000F | CCGGTACCTGAGCTCGCTAGCTGAATCCTAGAGAGGTGAGCTAGAGGAAAG |
| *Hnf4a* P-350F | CCGGTACCTGAGCTCGCTAGCCCACTCTCCCTCCCTCCTCT |
| *Hnf4a* P-0R | CAGTACCGGATTGCCAAGCTTTTATCTTATTGATTCTTCTAATCACCCAAGGTGG |

**Table S2. Antibodies used in the study**

| **IHC Staining** | | | | |
|---|---|---|---|---|
| Antibody | Species | Dilution | Company | Cat.No. |
| HNF4A | Rabbit | 1:200 | Cell Signaling | CST#3113 |
| CEBPA | Rabbit | 1:500 | Sigma-Aldrich | HPA065037 |
| p-SMAD2 | Rabbit | 1:100 | IBL | 28027 |
| GLUT2 | Rabbit | 1:300 | Sigma-Aldrich | HPA028997 |
| Anti-Rabbit/HRP | Swine | 1:200 | DAKO | P021702-2 |
| **Immunofluorescence** | | | | |
| E-Cadherin | Rabbit | 1:200 | Cell Signaling | CST#3195 |
| FOXO1 | Rabbit | 1:100 | Cell Signaling | CST#2880S |
| DRAQ5 | | 1:1000 | Cell Signaling | CST#4084 |
| Anti-Rabbit | Goat | 1:200 | ThermoFisher | A-21428 |

| **Immunoblotting** | | | |
|---|---|---|---|
| Antibody | Species | Company | Cat.No. |
| HNF4A | Rabbit | Cell Signaling | CST#3113 |
| CEBPA | Rabbit | Cell Signaling | CST#8178 |
| p-SMAD2 | Rabbit | Cell Signaling | CST#3108S |
| p-SMAD3 | Rabbit | Abcam | Ab63403 |
| SMAD2 | Rabbit | Cell Signaling | CST#5339S |
| SMAD3 | Rabbit | Cell Signaling | CST#9523S |

| | | | |
|---|---|---|---|
| p-AKT(Ser473) | Rabbit | Cell Signaling | CST#4060 |
| AKT | Rabbit | Cell Signaling | CST#4691 |
| H3K27ac | Rabbit | Abcam | Ab4729 |
| MED14 | Mouse | Santa Cruz | SC-81236 |
| E-Cadherin | Rabbit | Cell Signaling | CST#3195 |
| p-FOXO1(Ser256) | Rabbit | Cell Signaling | CST#9461 |
| GAPDH | Mouse | Santa Cruz | SC-25778 |
| β-ACTIN | Mouse | Santa Cruz | SC-47778 |
| Anti-Rabbit IgG | Mouse | Santa Cruz | SC-2357 |
| Anti-Mouse IgG | Goat | Santa Cruz | SC-2005 |
| **ChIP assay** | | | |
| H3K4me3 | Rabbit | Abcam | Ab8580 |
| RNA PoII Ser2 | Rabbit | Abcam | Ab193468 |
| RNA PoII Ser5 | Rabbit | Abcam | Ab5131 |
| Histone 3 | Rabbit | Abcam | Ab1791 |
| H3K27ac | Rabbit | Abcam | Ab4729 |
| CBP | Rabbit | Cell Signaling | CST#7389S |
| SMAD2 | Rabbit | Cell Signaling | CST#5339S |
| SMAD3 | Rabbit | Cell Signaling | CST#9523S |
| CEBPA | Rabbit | Invitrogen | PA5-77911 |
| FOXO1 | Rabbit | Cell Signaling | CST#2880S |
| IgG Control | Rabbit | Cell Signaling | CST#3900S |

**Table S3. siRNA and recombinant DNA used in the study**

| **Small interfering RNA** | | | |
|---|---|---|---|
| Product | Species | Company | Cat.No. |
| siCEBPA | Human | Santa Cruz | SC-37047 |
| siCebpa | Mouse | Santa Cruz | SC-37048 |
| siSMAD2 | Human | Thermo Fisher | 115714 |
| siSmad2 | Mouse | Thermo Fisher | 156216 |
| siSMAD3 | Human | Thermo Fisher | 107876 |
| siSmad3 | Mouse | Thermo Fisher | 156947 |
| siCREBBP | Human/Mouse | Thermo Fisher | 114409 |
| siTaf6 | Mouse | Thermo Fisher | 186593 |
| siTaf9 | Mouse | Thermo Fisher | 81626 |
| siCdk8 | Mouse | Thermo Fisher | 223930 |
| siMed14 | Mouse | Thermo Fisher | 70692 |
| siFoxo1 | Human/Mouse | Thermo Fisher | 106652 |
| Control siRNA | | Santa Cruz | SC-37007 |
| **Recombinant DNA** | | Company | Cat.No. |
| pFlag-CMV4-CEBPA | | Biomed | BM1310 |
| pFlag-CMV4 | | Biomed | |
| pScalps-mcebpa | | Addgene | 79551 |
| pScalps_Puro | | Addgene | 99636 |

| | | |
|---|---|---|
| pBabe RFP1-Smad2 neo | Addgene | 58491 |
| pBabe mRFP1 neo | Addgene | 98397 |
| mNeonGreen-Smad3 | Addgene | 172098 |
| mNeonGreen | Addgene | 189976 |
| α-RE | Provided by Atsushi Miyajima[1]. | |
| pGL4.14[luc2/Hygro] | Promega | E6691 |
| pGL4.74[hRluc/TK] | Promega | E6921 |

**Table S4. The sequences of siRNA**

| siRNA | **Sense (5'-3')** | **Antisense (5'-3')** |
|---|---|---|
| siCEBPA (sc-37047A) | CUCCAAUGCCUACUGAGUAtt | UACUCAGUAGGCAUUGGAGtt |
| siCEBPA (sc-37047B) | CCACGCCUGUCCUUAGAAAtt | UUUCUAAGGACAGGCGUGGtt |
| siCEBPA (sc-37047C) | CCGAUAUCAACACUUGUAUtt | AUACAAGUGUUGAUAUCGGtt |
| siCebpa (sc-37048A) | CGCUGGUGAUCAAACAAGAtt | UCUUGUUUGAUCACCAGCGtt |
| siCebpa (sc-37048B) | CCCACUUGCAGUUCCAGAUtt | AUCUGGAACUGCAAGUGGGtt |
| siCebpa (sc-37048C) | AGACCACCAUGCACCUACAtt | UGUAGGUGCAUGGUGGUCUtt |
| siSMAD2 (115714) | GGAGUGCGCUUAUACUACAtt | UGUAGUAUAAGCGCACUCCtc |
| siSmad2 (156216) | CGGUUAGAUGAGCUUGAGAtt | UCUCAAGCUCAUCUAACCGtc |
| siSMAD3 (107876) | GGACAGCGGGAAAAAUCGAtt | UCGAUUUUUCCCGCUGUCCtg |
| siSmad3 (156947) | GCGUAUAGGUGAUGUACAGtt | CUGUACAUCACCUAUACGCtg |
| siCREBBP (114409) | CGACAAAUCAUCUCUCAUUtt | AAUGAGAGAUGAUUUGUCGtg |
| siTaf9 (81626) | GGUUAAGUGUUGGUUCAGUtt | ACUGAACCAACACUUAACCtt |
| siTaf6 (186593) | CGACAGAAGCUCACUACCAtt | UGGUAGUGAGCUUCUGUCGtt |
| siMed14 (70692) | GGAAAUCUGAUGUGGAAAGtt | CUUUCCACAUCAGAUUUCCtt |
| siCdk8 (223930) | CCACAUCUUCACAAUGUCCtt | GGACAUUGUGAAGAUGUGGtt |
| siFoxo1 (106652) | GGCAUCUCAUAACAAAAUGtt | CAUUUUGUUAUGAGAUGCCtg |

**Table S5. Reagent kits used in the study**

| Critical Commercial Assays | | |
|---|---|---|
| Product | Company | Cat.No. |
| Lipofectamine 3000 kit | Thermo Fischer | L3000008 |
| Lipofectamine® RNAiMAX kit | Thermo Fischer | 13778-075 |
| PureLink Quick Plasmid Miniprep Kit | Invitrogen | K210010 |
| PureLink® HiPure Plasmid Maxiprep Kit | Invitrogen | K210007 |
| Phusion® High-Fidelity DNA Polymerase | New England Biolabs | M0530S |
| Gibson Assembly HiFi Cloning Kit | Thermo Fisher | A46624 |
| SYBR Green Master Kit | Thermo Fisher | A25918 |
| Glucose uptake assay | Promega | J1341 |
| Luciferase assay system | Promega | E1500 |
| Dual-Luciferase Assay Kit | Promega | E2920 |
| Human Insulin ELISA KIT | Sigma-Aldrich | RAB0327 |
| Mouse Insulin ELISA KIT | Thermo Fischer | EMINS |

Supplementary figures and figure legends

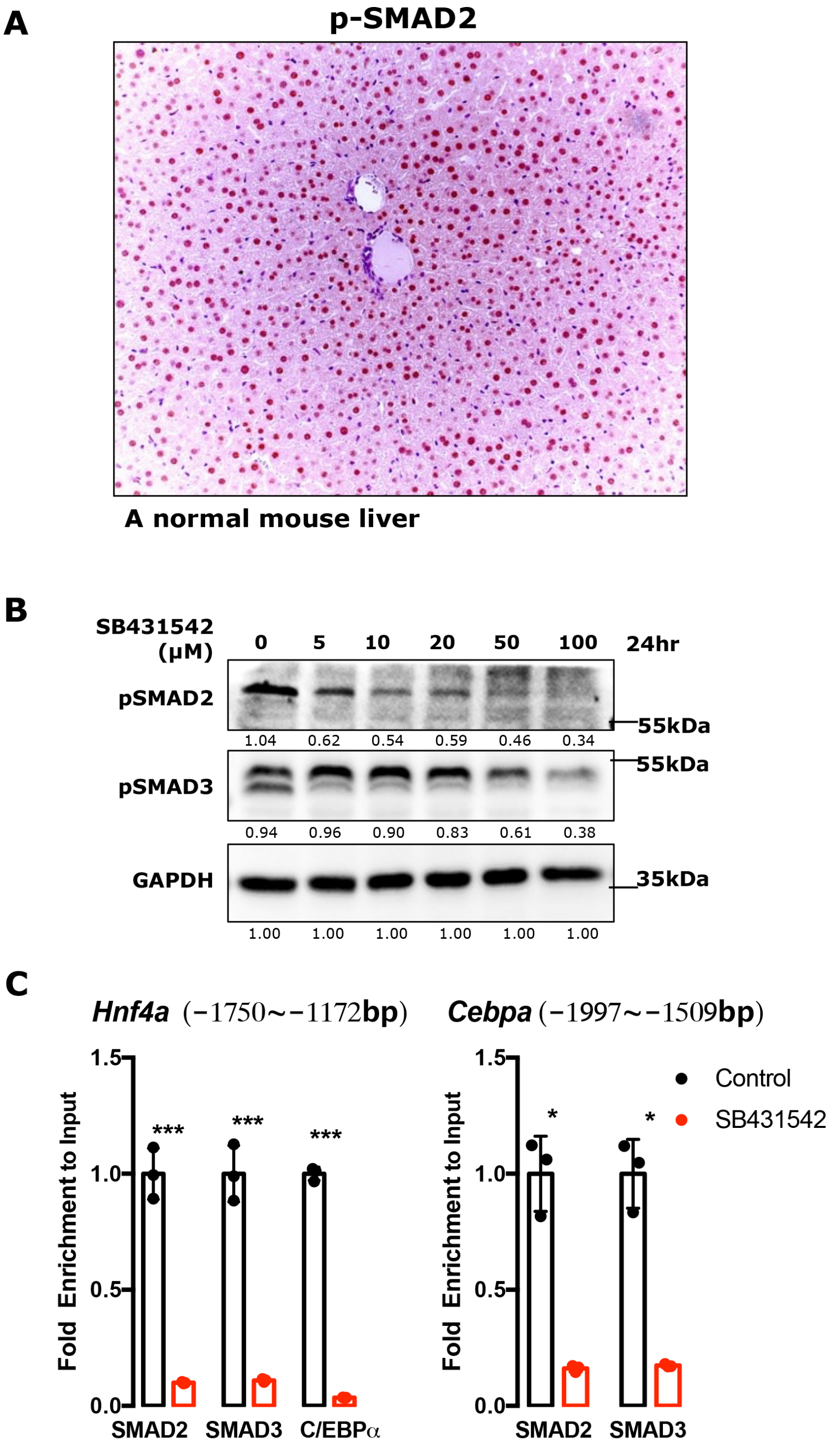


**Figure S1. The existence of activated SMAD proteins in normal mouse livers and cultured hepatocytes.** (A) Immunohistochemical staining for p-SMAD2 was performed in 20 normal mouse liver tissues. A representative picture depicts p-SMAD2

staining in hepatocytes. **(B)** Western blotting shows the effects of different doses of TGF-β receptor I inhibitor SB431542 incubation for 24 h on p-SMAD2 and p-SMAD3 expression in AML12 cells. GAPDH was used as loading control. **(C)** ChIP assays were performed to examine the binding of SMAD2 or SMAD3 to the *Hnf4a* and *Cebpa* promoters in AML12 cells with or without SB431542 treatment. Bars represent the mean ± SD, P-values were calculated by One-way ANOVA, *: $P<0.05$; **: $P<0.01$; ***: $P<0.001$. Triple experiments were performed, and one representative experiment is shown.

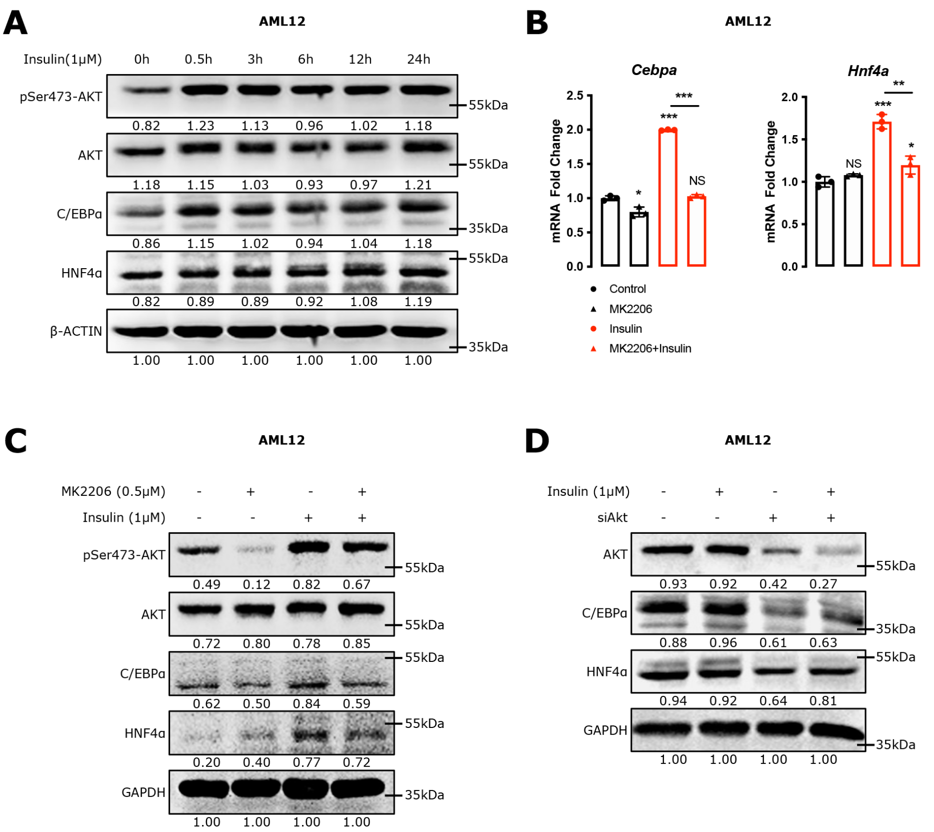


**Figure S2. Insulin is crucial for the maintenance of C/EBPα expression in hepatocytes.**

**(A)** Western blotting analyzed the effect of dynamic insulin (1 μM) treatment on C/EBPα and HNF4α expression in AML12 cells. **(B-C)** qPCR **(B)** and western blotting **(C)** analyzed C/EBPα and HNF4α expression in AML12 cells. The cells were treated with pan-AKT inhibitor MK2206 (0.5 μM) for 1 hour followed by insulin (1 μM) stimulation for 24 hours. **(D)** Western blotting examined C/EBPα and HNF4α expression in AML12 cells. The cells were treated with *Akt*-siRNA for 24 hours, subsequently stimulation with insulin (1 μM) for another 24 hours. Bars represent the mean ± SD, *P*-values were calculated by One-way ANOVA, *: *P*<0.05; **: *P*<0.01; ***: *P*<0.001; and NS, no significance. Triple experiments were performed, and one representative experiment is shown.
